# Basis of executive functions in fine-grained architecture of cortical and subcortical human brain networks

**DOI:** 10.1101/2022.12.01.518720

**Authors:** Moataz Assem, Sneha Shashidhara, Matthew F. Glasser, John Duncan

**Author notes:** **Corresponding author**; Address: 15 Chaucer Road, Cambridge, UK, CB2 7EF.

## Abstract

Theoretical models suggest that executive functions rely on both domain-general and domain-specific processes. Supporting this view, prior brain imaging studies have revealed that executive activations converge and diverge within broadly characterized brain networks. However, the lack of precise anatomical mappings has impeded our understanding of the interplay between domain-general and domain-specific processes. To address this challenge, we used the high-resolution multimodal MRI approach of the Human Connectome Project to scan participants performing three canonical executive tasks: n-back, rule switching, and stop signal. The results reveal that, at the individual level, different executive activations converge within 9 domain-general territories distributed in frontal, parietal and temporal cortices. Each task exhibits a unique topography characterized by finely detailed activation gradients within domain-general territory shifted towards adjacent resting-state networks; n-back activations shift towards the default mode, rule switching towards dorsal attention and stop signal towards cingulo-opercular networks. Importantly, the strongest activations arise at multimodal neurobiological definitions of network borders. Matching results are seen in circumscribed regions of the caudate nucleus, thalamus and cerebellum. The shifting peaks of local gradients at the intersection of task-specific networks provide a novel mechanistic insight into how partially-specialised networks interact with neighbouring domain-general territories to generate distinct executive functions.

## Introduction

Executive function is an umbrella term for the processes necessary to manage diverse cognitive challenges. The range of “executive tasks” is vast and includes recalling or manipulating items in short-term memory, generating verbs under time pressure, withholding a habitual motor response, attending to specific stimuli and ignoring distractors, solving tasks with constantly changing rules, and following complex instructions (Miyake et al. 2000; Diamond 2013). Performance on executive tasks can identify severe cognitive deficits in patients with brain lesions, correlates with measures of general intelligence and predicts real life problem solving abilities (Duncan et al. 1996; Miyake et al. 2000; Roca et al. 2010; Friedman and Miyake 2017; Woolgar et al. 2018). Despite much progress, the underpinning of executive processes in the brain is still only partly understood.

One approach to identify executive processes examines individual differences in executive task performance. An influential “unity and diversity” model found that performance on all executive tasks tends to positively correlate, suggesting a common underlying process usually referred to as the “common executive function” (Miyake et al. 2000; Friedman and Miyake 2017) and linked to several constructs in theoretical models such as the central executive in working memory (Baddeley and Hitch 1974), the *g*-factor for general intelligence (Spearman 1904), proactive control in the dual mechanisms framework (Braver 2012) and energization (Stuss and Alexander 2000). To explain the remaining variance, the model also proposes several more specialized components, with the original model (Miyake et al. 2000) identifying three putative processes labelled as updating, set shifting and inhibition. Recent replications have highlighted that fine-scaled division of components varies with diversity of the task battery, model chosen and the age of participants (Karr et al. 2018). Theoretical models thus suggest the existence of both domain-general and domain-specific brain processes to support executive task performance.

Another approach concerns brain lesion studies, the historical driver for the development of executive tasks. Relatively circumscribed lesions in frontal and parietal cortices are associated with widespread deficits in executive performance (Roca et al. 2010; Woolgar et al. 2010, 2018), suggesting a domain-general process that has been compromised. A finer-grained view of lesion data, however, has often been used to argue that distinct executive functions are supported by distinct frontal lobe territories. For example, an authoritative review of two decades of brain lesion studies concluded that “there is no central executive”. Instead, it attributed distinct executive processes to distinct territories: energization (dorsomedial), monitoring (right lateral), task setting (left lateral), emotional self-regulation (ventromedial) and metacognition (fronto-polar) (Stuss 2011). The spatially coarse nature of human brain lesions has hindered our ability to provide a comprehensive neurobiological explanation for the interplay between domain-general and domain-specific processes.

Functional magnetic resonance imaging (fMRI) studies in healthy participants have provided a more detailed picture. On the one hand, meta-analysis and within-subject studies of diverse executive functions show circumscribed overlaps in the lateral and dorso-medial frontal cortices, insula, intraparietal sulcus and occipito-temporal junction (Collette et al. 2005, 2006; Niendam et al. 2012; Fedorenko et al. 2013; Nee et al. 2013; Lemire-Rodger et al. 2019; Braver et al. 2021; He et al. 2021; Saylik et al. 2022; Friedman and Robbins 2022; Reineberg et al. 2022). These activations are usually linked to domain-general or multiple-demand (MD) areas that co-activate in association with many cognitively demanding tasks (Duncan 2010; Fedorenko et al. 2013; Assem et al. 2020; Shashidhara et al. 2020). On the other hand, several studies have fractionated activations based on the statistical strength of their engagements in different executive processes (Wager et al. 2005; Dosenbach et al. 2006; Dodds et al. 2011; Hampshire et al. 2012; Niendam et al. 2012; Lemire-Rodger et al. 2019; He et al. 2021; Reineberg et al. 2022). While such results are broadly in line with a picture of both unity and diversity, as yet there is no clear consensus on how domain-general and domain-specific executive processes combine.

Our recent work using high quality multi-modal imaging approaches of the Human Connectome Project (HCP) suggests a clearer path forward for investigating the putative unity and diversity of executive functions. HCP methods utilize surface-based approaches and multimodal MRI features for accurate alignment of cortical regions across individuals (Glasser, Coalson, et al. 2016; Glasser, Smith, et al. 2016). Previously we used HCP data to refine the anatomy of MD activations, delineating 9 MD cortical patches per hemisphere distributed in frontal, parietal and temporal lobes (Assem et al. 2020) (**Figure 1a**). Within the 9 patches, using the HCP’s recent multimodal cortical parcellation (HCP MMP1.0), we defined an MD core consisting of 10 out of 180 MMP1.0 areas per hemisphere, that are most strongly co-activated across multiple task contrasts, and most strongly functionally interconnected, surrounded by a penumbra of 18 additional regions [**Figure 1b**; (Assem et al. 2020)]. This fine-grained picture of the MD system highlights several challenges for interpreting previous executive function studies. First, while executive activations often appear to overlap MD regions (Friedman and Miyake 2017), it remains unknown whether executive tasks engage penumbra or core MD regions, or additional nearby regions with more task-specific responses. Second, links between executive activations and canonical resting state networks (RSNs) are uncertain. Previous studies propose overlaps with the fronto-parietal network (FPN) (Power et al. 2011; Yeo et al. 2011; Blank et al. 2014; Ji et al. 2019; Assem et al. 2020; Cocuzza et al. 2020). In our study we used the RSN definitions from Ji et al. (2019), which are based on HCP resting state data and the Glasser et al. (2016) parcellation. We found that core MD regions formed a functionally integrated subset of the Ji et al. (2019) FPN. Penumbra MD regions included further FPN regions, along with regions from three other RSNs, the cingulo-opercular network (CON), the dorsal attention network (DAN) and the default mode network (DMN). We have hypothesized that such nearby nodes could act as communication channels between domain-specific and MD regions. It is currently unclear how different executive activations relate to RSNs. Third, previous studies have largely ignored subcortical and cerebellar contributions to executive functions (Niendam et al. 2012). Our previous work identified circumscribed MD regions in the head of caudate and localized patches in cruses I and II in the cerebellum as well as a putative MD region in the anterior and medial thalamus (Assem et al. 2020, 2022). The relation between non-cortical MD regions and executive activations remains uncharted territory.

**Figure 1.**
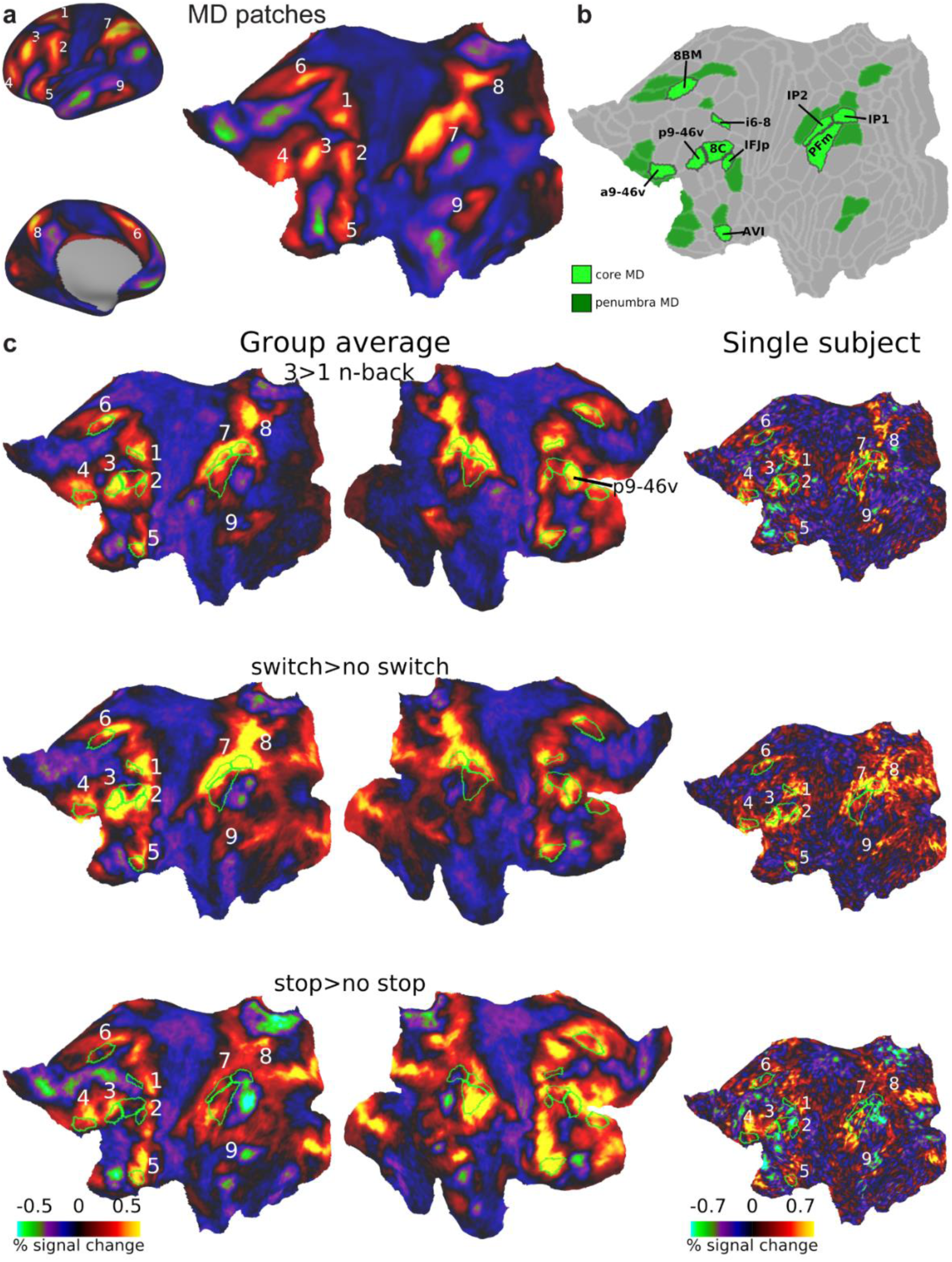
**(a)** The nine MD patches displayed on cortical surface (left) and a flattened left surface (right) as revealed by average activations of 449 subjects based on three cognitively demanding contrasts from (Assem et al. 2020): 2>0 n-back, hard>easy reasoning, math>story. **(b)** Extended MD system from (Assem et al. 2020). Core MD regions are colored in bright green surrounded by black borders and individually labelled. Penumbra MD regions are colored in dark green. Data available at: http://balsa.wustl.edu/r27NL **(c)** Flat cortical maps overlaid with group average activations for each executive contrast in the current study. Green borders surround core MD areas, with the nine coarser-scale patches labelled on the left hemisphere. Right column shows example activation from a single subject on the left hemisphere. See Supplementary Figure 1 for more single subject activations. All single subject data is available at: http://balsa.wustl.edu/x8M0q

To investigate executive activations and their relation to MD regions with high spatial precision, we collected a new dataset using HCP’s multimodal MRI acquisition and analysis approach. We chose three classical paradigms targeting three putative executive functions: an n-back task (updating), a rule switching task (set shifting) and a stop signal task (inhibition). The same subjects performed all three tasks within the same session and within the same runs. In all three cases, a high-demand executive condition was contrasted with a low-demand baseline. This is a critical manipulation because MD regions are characterized by their strong response to task difficulty manipulations (Fedorenko et al. 2013; Assem et al. 2020).

Our results explicate both unity and diversity of executive functions. The resulting scheme, however is quite different from classical views of distinct frontal territories and provides new mechanistic insights interlinking domain-general and domain-specific processes. The results show that the three executive tasks show overlapping activations at the single subject level within MD patches, suggesting a common role for MD regions in executive tasks. Yet each task’s topography shifts within MD patches to form a unique intersection between core MD and adjacent fine-grained RSNs. In this intersection, the strongest activations often arise at the border between a core MD region and an adjacent RSN. These results suggest a novel framework for how domain-specific areas recruit neighboring MD areas to generate distinct executive functions. They provide a new, fine-scale resolution of longstanding debates between domain-specific and domain-general views of executive function.

## Materials and Methods

### Subjects

Thirty-seven human subjects participated in this study (age=25.9±4.7, 23 females, all right-handed). Originally fifty subjects were scanned over two sessions; thirteen subjects were excluded either due to incomplete data (n=5), excessive head movement during scanning (n=4; movement more than double the fMRI voxel size), technical problems during scanning (n=2; MRI scanner crashing) or during analysis (n=2; excessive field inhomogeneities due to unreported teeth implants that affected structural scans). All subjects had normal or corrected vision (using MRI compatible glasses). Informed consent was obtained from each subject and the study was approved by the Cambridge Psychology Research Ethics Committee.

### Task Paradigms

Each subject performed three tasks in the same scanning session: n-back, switch and stop signal (**Figure 2**). All three tasks were visual. Subjects underwent a resting-state scan in a second session. Before scanning, participants performed a short training session ensuring they understood the instructions and were performing above chance. This is particularly important for the stop signal task (see below).

**Figure 2.**
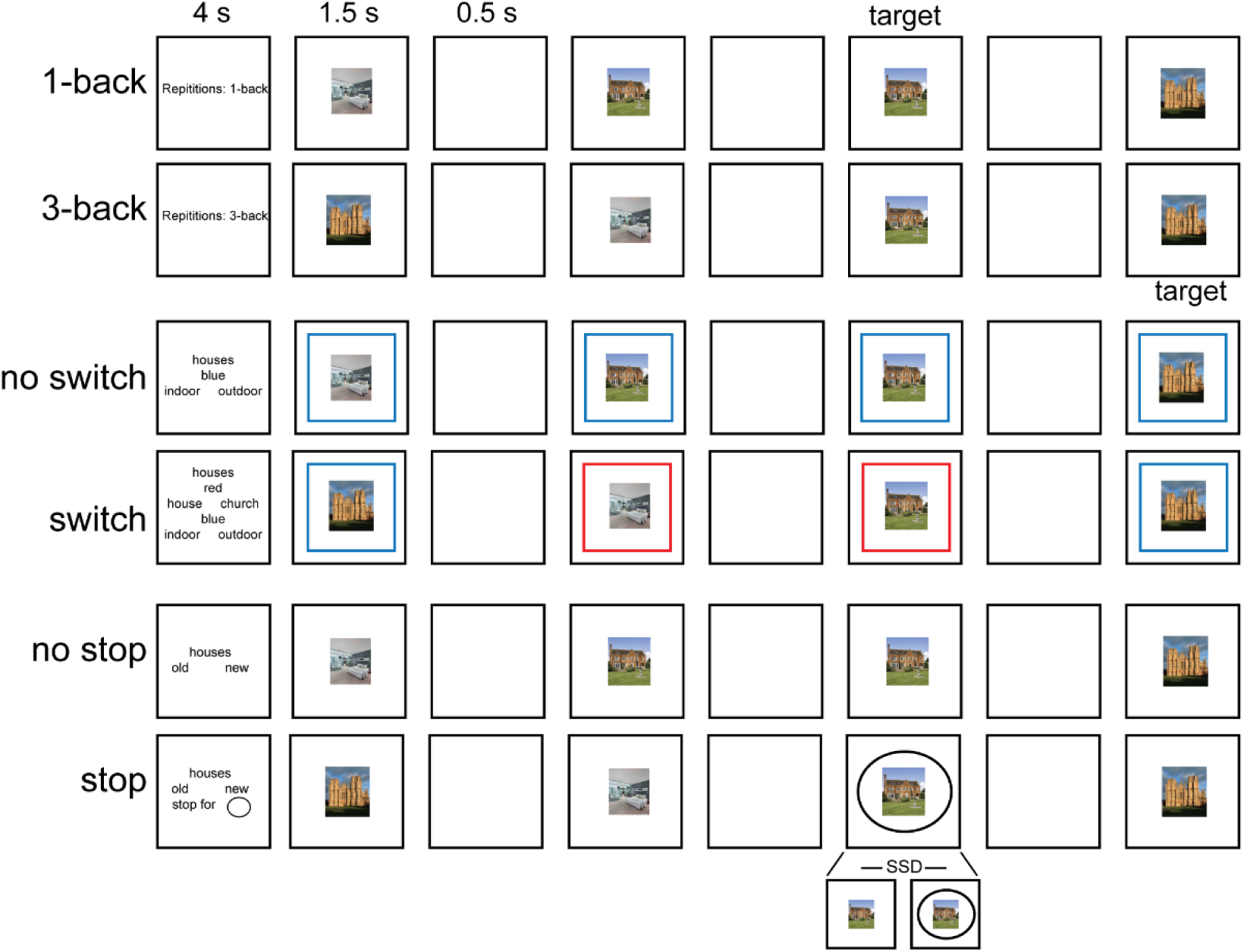
Illustration of the three tasks performed in the current study. Note stimuli were either faces or houses.

Each subject performed four runs. Each run consisted of 36 blocks: 8 n-back, 8 switch, 8 stop and 12 fixation blocks. Each task consisted of 4 easy and 4 hard blocks. Each task block (30 s) started with a cue (4 s) followed by 12 trials (24 s, 2 s each) and ended with a blank screen (2 s) as an inter-block interval. Easy and hard blocks of one task were paired (easy followed by hard, or hard followed by easy) and the order was counterbalanced across runs and subjects. A fixation block (16 s) followed every two paired task blocks. For each trial in the task blocks, the visual stimulus was presented for 1500 ms, followed by 500 ms of a blank screen. Responses were accepted at any moment throughout the trial. Stimuli were pictures of faces or houses (each category in a separate block). Face stimuli were selected from the Developmental Emotional Faces Stimulus Set (Meuwissen et al. 2017). Faces were either males or females, children or adults, making a happy or sad face. House stimuli were pictures of houses or churches, old or new, from inside or outside. There were 32 faces and 32 houses, each made up of 4 examples for each of the 2 x 2 x 2 possible feature combinations. Subjects were encouraged to use their right hand and respond to targets using a middle finger press and to non-targets using an index finger press but this was not enforced and several subjects found it more comfortable to use both hands for responses (index fingers or thumbs).

#### N-back task

For the 3-back condition (hard), subjects were instructed to press right for the target stimulus (i.e. current stimulus was the same as the one 3 steps back), and left for all non-target presentations. Similarly, for the 1-back condition (easy), subjects were instructed to press right for the target stimulus (i.e. current stimulus was an exact repetition of the immediate previous stimulus) and press left for all non-target stimuli. In each block there were 1-2 targets and 2 lures (a target image but at the 2-back or 4-back positions).

#### Switch task

The switch rules were indicated by colored screen borders. The colors were either red or blue. For the 1-rule blocks (easy), the border color did not change throughout the trials of a single block. If the stimuli were faces, a red border indicated to the participant to respond whether the face was male (left press) or female (right press), while a blue border required a judgement if the face was that of a child (left press) or an adult (right press). If the stimuli were houses, for a red border participant responded whether the house was a standard house (left press) or a church (right press), while a blue border required a judgement if the picture was indoor (left press) or outdoor (right press). For the 2-rule blocks (hard), the colored borders would change randomly throughout the trials of a single block, ensuring an equal number of red/blue borders per block.

#### Stop signal task

For the no stop blocks, participants pressed left if the stimulus was a happy face (or old house) and pressed right if it was a sad face (or a new house). For the stop blocks, 33% of trials were stop signal trials. The instructions for the stop block were the same except that, if a stop signal appeared (a black circle surrounding the central stimulus), participants were instructed to withhold their response. To discourage participants from responding slowly, we employed a tracking procedure for the stop signal delay (SSD), i.e. the delay between onset of the face/house stimulus and the black circle. On each stop signal trial, SSD was set to 200 ms below the running average of the subject’s reaction time (RT) on all previous go trials for the same stimulus category, including practice trials.

### Image Acquisition

Images were acquired using a 3T Siemens Prisma scanner with a 32-channel RF receive head coil. MRI CCF acquisition protocols for HCP Young Adult cohort were used (package date 2016.07.14; https://protocols.humanconnectome.org/CCF/). These protocols are substantially similar to those described in previous studies (Glasser et al. 2013; Smith et al. 2013; Uǧurbil et al. 2013) but do differ in some respects. All subjects underwent the following scans over two sessions: structural (at least one 3D T1w MPRAGE and one 3D T2w SPACE scan at 0.8-mm isotropic resolution), rest fMRI (2 runs × 15 min), and task fMRI (3 tasks, 4 runs each, approx. 60 min total). Whole-brain rest and task fMRI data were acquired using identical multi-band (factor 8) gradient echo EPI sequence parameters of 2-mm isotropic resolution (TR = 800 ms, TE=37 ms). Both rest and task EPI runs were acquired in pairs of reversed phase-encoding directions (AP/PA). Spin echo phase reversed images (AP/PA) matched to the gradient echo fMRI images were acquired during the structural and functional (after every 2 functional runs) scanning sessions to (1) correct T1w and T2w images for readout distortion to enable accurate T1w to T2w registration, (2) enable accurate cross-modal registrations of the fMRI images to the T1w image in each subject, (3) compute a more accurate fMRI bias field correction and (4) segment regions of gradient echo signal loss.

### Data preprocessing

Data preprocessing was also substantially similar to the HCP’s minimal preprocessing pipelines detailed previously (Glasser et al. 2013). A brief overview and differences are noted here. HCP pipelines versions 3.27.0 were used (scripts available at: https://github.com/Washington-University/HCPpipelines). For each subject, structural images (T1w and T2w) were used for extraction of cortical surfaces and segmentation of subcortical structures. Functional images (rest and task) were mapped from volume to surface space and combined with subcortical data in volume to form the standard CIFTI grayordinates space. Data were smoothed by a 2mm FWHM kernel in the grayordinate space that avoids mixing data across gyral banks for surface data and avoids mixing across major structure borders for subcortical data.

From this point onwards HCP pipelines version 4.0.0 were used (also available through the link above; specific parameters different from the default values are noted below). Rest and task fMRI data were additionally identically cleaned up for spatially specific noise using spatial ICA+FIX (Salimi-Khorshidi et al. 2014). ICA+FIX was applied separately to each of the following concatenated runs: resting-state runs (2x15 mins), task runs from session one (4x15 mins). An improved FIX classifier was used (HCP_Style_Single_Multirun_Dedrift in ICAFIX training folder) for more accurate classification of noise components in task fMRI datasets. After manual checking of ICA+FIX outputs for 10 subjects, a threshold of 50 was determined for “good” vs “bad” signal classification and applied for the remaining subjects. In contrast to the Assem et al. (2020) study, global structured noise, largely from respiration, was not removed using temporal ICA as public scripts were not yet publicly available at the time the data were analyzed.

For accurate cross-subject registration of cortical surfaces, the multimodal surface matching algorithm MSM was used. First “sulc” cortical folding maps are gently registered in the MSMSulc registration, optimizing for functional alignment without overfitting folds. Second, a combination of myelin, resting-state network, and rest fMRI visuotopic maps (Robinson et al. 2014, 2018) is used to fully functionally align the data. For this purpose we used the 30 mins of resting state data.

### Task fMRI analysis

Task fMRI analysis scripts in HCP pipelines version 4.0.0 were used. Default steps are detailed in Barch et al. (2013). Briefly, autocorrelation was estimated using FSL’s FILM on the surface (default parameters in HCP’s task fMRI analysis scripts were used). Activation estimates were computed for the preprocessed functional time series from each run using a general linear model (GLM) implemented in FSL’s FILM (Woolrich et al. 2001).

For each of the tasks, 4 regressors were used (2 stimulus category x 2 task difficulty). Each predictor had a unitary height and covered the period from the onset of the cue to the offset of the final trial (28 sec). All regressors were then convolved with a canonical hemodynamic response function and its temporal derivative. 12 additional motion regressors were added to the model (3 translation, 3 rotation and their derivatives). The time series and the GLM design were temporally filtered with a Gaussian-weighted linear highpass filter with a cutoff of 200 seconds. Finally, the time series was prewhitened within FILM to correct for autocorrelations in the fMRI data. Surface-based autocorrelation estimate smoothing was incorporated into FSL’s FILM at a sigma of 5mm. Fixed-effects analyses were conducted using FSL’s FEAT to estimate the average effects across runs within each subject.

For further analysis of effect sizes, beta ‘cope’ maps were moved from the CIFTI file format to the MATLAB workspace. Beta maps were then converted to percent signal change as follows: 100*(beta/10000). The value 10000 corresponds to the mean scaling of each vertex/voxel’s timeseries during preprocessing. Unless mentioned otherwise, parametric statistical tests were used.

For parcellating the cerebral cortex, the group-average HCP multi-modal parcellation (MMP1.0) was used (Glasser, Coalson, et al. 2016), as the individual-specific areal classifier is not publicly available. Still, due to the superior cortical alignment approach of MSMAll, the areal fraction of individually defined parcels captured by group-defined borders reaches 60–70% (Coalson et al. 2018) and we have previously demonstrated that using areal classifier and group-defined borders produces similar results (Assem et al. 2020). Where appropriate, values of vertices sharing the same areal label were averaged together to obtain a single value for each area.

To create the RGB colors in **Figure 5a**, we converted each task’s group average activation map to lie between 0 and 1 by normalizing their activations by the minimum and maximum activation value across all three contrasts as follows: (value-OverallMinValue)/(OverallMaxValue-OverallMinValue). Each vertex was then assigned a color through a 1x3 vector [red green blue] with the value of each color ranging from 0 to 255. The color was assigned by combining the normalized activations of the three tasks as follows [n-back switch stop]*255.

### Resting-state connectivity analysis

For connectivity analysis we used a dense connectivity matrix (59k by 59k vertices) from 210 HCP subjects [the 210 validation group from (Glasser, Coalson, et al. 2016)]. Each subject underwent 1 hour of resting-state scans. The analysis methods are described in detail in (Glasser, Coalson, et al. 2016). Briefly, the pipeline was very similar to this current study with the addition of temporal ICA cleaning to remove global respiratory artefacts (Glasser et al. 2018, 2019).

### Borders analysis and simulation

We first calculated the geodesic distance between all cortical vertices using the connectome workbench function -surface-geodesic-distance-all-to-all using subject-specific vertex areas and the midthickness cortical surface (derived from our 37 subjects). We then identified the vertices that belonged to the HCP_MMP1.0 borders using the workbench function –border-to-vertices. Vertices with each area were then sorted according to their distance from the border vertices and grouped into 5 distance groups by ensuring a similar number of vertices was included across all groups. Border vertices were included in this analysis.

To create the simulated data, we randomly selected 37 subjects from the 449 HCP subjects with each subject’s cortex parcellated into 360 areas using a multimodal areal classifier (Glasser, Coalson, et al. 2016). We then populated the vertices for each area and each subject with activation values derived from our 37 subjects; e.g. if the n-back average activation value for subject_1 for area p9-46v was 0.3, we populated the vertices belonging to area p9-46v in HCP_subject_1 with 0.3. We created three simulated datasets corresponding to the three executive contrasts. We then applied smoothing for each subject using workbench’s function –cifti-smoothing with varying smoothing levels: 4, 12 and 20 mm FWHM to simulate inherent smoothness/noise of fMRI signal at multiple levels

### Subcortical and cerebellar analysis

The HCP minimal preprocessing pipeline utilizes FreeSurfer’s standard segmentation carried out separately for every subject. The 19 subcortical/cerebellar structures are left and right caudate, putamen, globus pallidus, thalamus, cerebellum, hippocampus, amygdala, ventral diencephalon, nucleus accumbens; plus whole brain stem. In this study we focused on the caudate, thalamus and cerebellum.

For the voxel-wise conjunction analysis in caudate and thalamus, we applied an additional 4 mm FWHM to the data using the workbench function –cifti-smoothing. All other analyses in this section, including all analyses of cerebellar activation, used unsmoothed data.

In Assem et al (2020) two versions of the subcortical/cerebellar MD masks were defined: One based on a conjunction of task activations and one based on rfMRI connectivity with cortical MD core. In the current study, the caudate and cerebellar masks were based on task activations as they are slightly more spatially constrained than the rfMRI mask. The thalamic MD mask was based on rfMRI, as our previous study could not identify a task-based conjunction in the thalamus. The volumetric cerebellar results are projected on a flat cerebellar surface using SUIT software (Diedrichsen and Zotow 2015). Although this approach has the limitations of a volume-based analysis (and thus is done mainly to aid visualization), individual subject cerebellar surface reconstruction and registration is not yet easily available.

## Results

50 subjects were scanned while performing three classical executive tasks in the same session: n-back, switch and stop signal. Data from 37 subjects were included in this report (see **Materials and Methods** for subjects’ details). All tasks were visually presented and required button presses. Each task had two difficulty conditions. The n-back task consisted of 3-back and 1-back blocks. The switch task had two rule and one rule blocks. The stop signal task had blocks with stop trials and blocks with no stop trials. Participants performed 4 runs, each lasting 15 minutes and containing 4 easy and 4 hard blocks for each task, along with 12 fixation blocks. Additionally each subject underwent 30 minutes of resting state scans in a separate session. (**Figure 2**; see **Materials and Methods** for further task details).

### Behavior

As expected, performance on the easy condition was better than the hard condition for all tasks (**Table 1**). For the targets in the n-back task, accuracy was higher and reaction times shorter for the 1-back condition than the 3-back condition (accuracy t_36_=6.6; reaction time (RT) t_36_=13.8, both *p*s<0.0001). Similarly, accuracy and RTs on the switch blocks were worse than the no switch blocks (accuracy t_36_=9.3; RT t_36_=27.2, both *p*s<0.0001). For the stop signal task, participants had more go omissions in the stop blocks (t_36_=3.0, *p*<0.01). In the stop block, participants successfully stopped on 44.7% (±10.6) of stop trials, with unsuccessful stop trial RTs faster than go trial RTs (**Table 1**).

**Table 1.** Performance on the three tasks [mean ± standard deviation].

| **N-back** | **1-back** | **3-back** |
| --- | --- | --- |
| Target accuracy (%) | 92.5±9.2 | 78.8±10.6 |
| Target RT (ms) | 623±80.6 | 819.5±110.5 |
| **Switch** | **No switch** | **Switch** |
| Accuracy (%) | 97.2±2.1 | 92.0±4.6 |
| RT (ms) | 725.5±90.2 | 985.6±90.3 |
| **Stop signal** | **No stop** | **Stop** |
| Go omission (%) | 0.3±0.6 | 1.0±1.9 |
| Go accuracy (%) | 96.1 ± 2.7 | 95.7±3.6 |
| Successful stops (%) | n/a | 44.7±10.6 |
| Unsuccessful stop RT (ms) | n/a | 932.8±240.8 |
| Correct Go RT (ms) | 722.2 ± 74.4 | 1063.0±269.7 |
| Stop signal delay (SSD; ms) | n/a | 855.1±269.8 |

### Overview of executive task activations

Structural and functional MRI data were preprocessed using surface-based approaches according to the HCP minimal preprocessing pipelines (see **Materials and Methods**). Additionally, functional data were cleaned using spatial ICA+FIX and were aligned across subjects using the multimodal surface matching algorithms utilizing cortical curvature, myelin and functional connectivity maps (MSMAll; see **Materials and Methods**). Data were not smoothed beyond the surface-based 2 mm FWHM smoothing in the preprocessing step.

We first sought an overview of activations for each of the three critical contrasts (3>1 n-back, switch>no switch, stop>no stop). **Figure 1c** reveals broad similarities between the three tasks, with activations resembling the 9 MD patches from Assem et al. (2020), within and also adjacent to the 10 finer-scale core regions (green borders). A partial exception is the temporal patch from Assem et al. (2020); though activations close to this patch were seen for all three contrasts (compare **Figure 1a**), they did not clearly overlap. Of note is the replicability of the 9 patches at the single subject level (**Figure 1c** and **Supplementary Figure 1)** The data also suggest that, in more detail, each task shows a unique pattern of activations. At the hemispheric level, there were stronger left hemispheric activations for switch, but right biased activations for stop. Within each hemisphere, exact activation patterns differed within and adjacent to MD regions. At a sub-areal finer grained level, for example, note activation patterns at the different edges of the right lateral prefrontal region p9-46v, with stop activations near its anterior-dorsal border, and switch activations near its posterior-ventral border.

With this broad overview, it seems plausible that executive activations harbor both similarities and differences. In the next sections we explore the similarities, differences and fine-grained activations across these three tasks.

### Executive activations converge on a common MD core at the individual level

First, we sought to statistically investigate conjunctions between the three executive contrasts at the coarse areal level. For areal definitions, we used the HCP’s MMP1.0 (Glasser, Coalson, et al. 2016). With the improved MSMAll alignment, previous work has shown that the HCP_MMP1.0 group-defined borders capture around 70% of the areal fraction of individually defined areas (Coalson et al. 2018) and produce closely matched results to those derived from subject-specific areal definitions (Assem et al. 2020, Assem et al 2022). For each contrast we identified the significantly activated areas across subjects (one sample t-test against zero, p<0.05 Bonferroni corrected for 360 cortical areas). A set of 31 areas showed a conjunction of all three significant contrasts in at least one hemisphere. The areas were spread throughout 8 of the 9 coarse-scale MD patches as there were no surviving areas in the temporal lobe (all 31 areas displayed on the left hemisphere in **Figure 3a**). At the finer scale of individual HCP regions, except for IP1, all remaining 9 core MD areas were co-activated in each contrast in at least one hemisphere, further confirming their domain-generality (**Figure 3a**). The remaining 22 regions included 10 of the 18 penumbra regions defined in Assem et al. (2020), which here we term 2020-penumbra, along with 12 new regions that we call additional-penumbra. On the dorso-lateral frontal surface, we identify new areas FEF, 6a and 6ma near core MD region i6-8. More anterior, we identify areas 46 and 9-46d near core MD areas p9-46v and a9-46v. In the inferior frontal junction we identify PEF abutting core MD region IFJp. In the insular region, we identify area FOP4 next to core MD region AVI. On the medial parietal surface, we identify a cluster of four areas next to POS2 (penumbra MD); these are 7Pm, 7PL, 7Am and PCV. All additional-penumbra regions were close to core MD regions.

**Figure 3.**
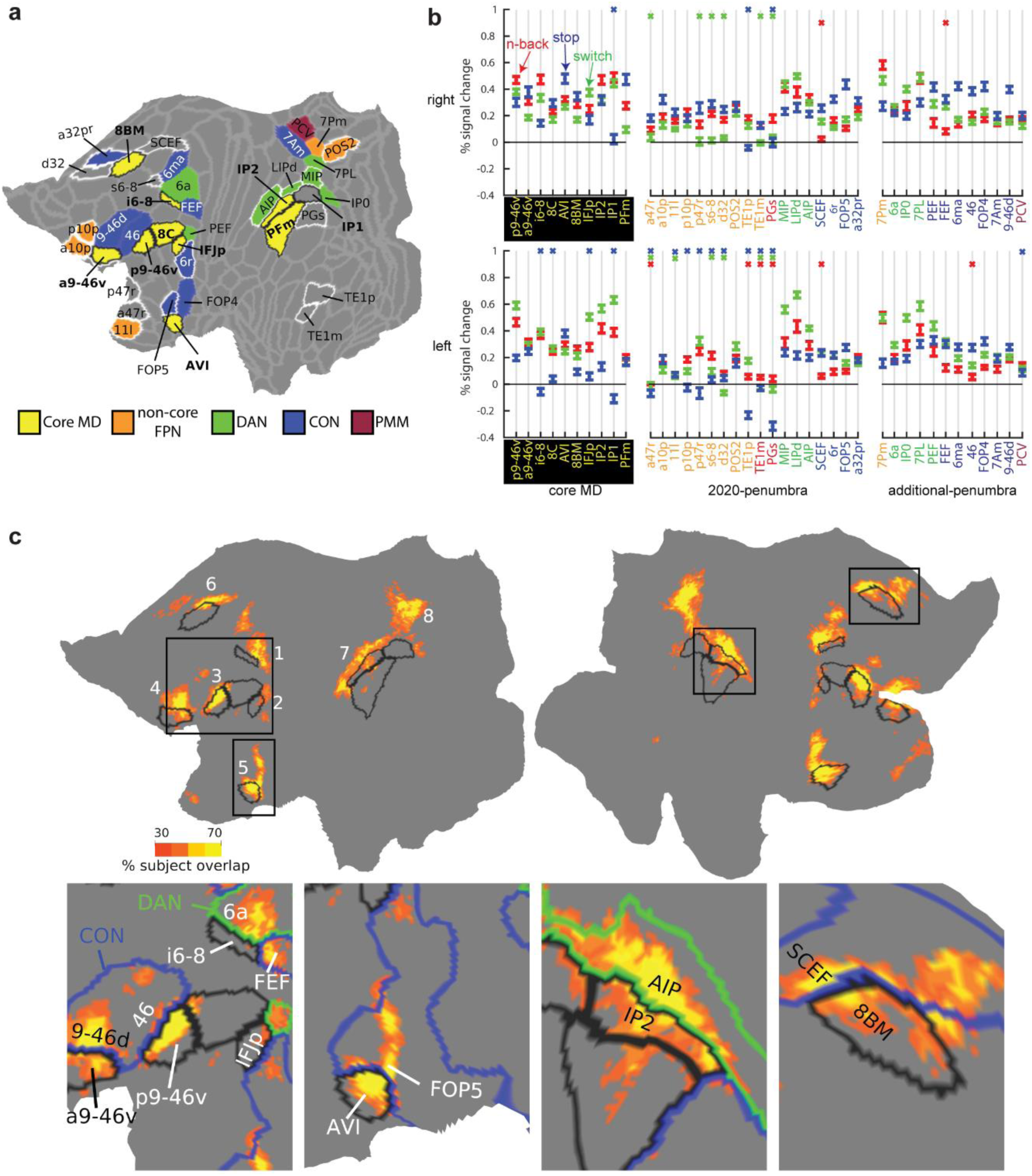
The unity of executive functions. **(a)** Cortical parcels showing conjunction of significant activation in each of the three executive contrasts (p<0.05 Bonferroni corrected). All unique areas identified in either hemisphere are projected on the left hemisphere, and colored according to RSN membership from Ji et al. (2019). Black borders surround core MD, white borders surround 2020-penumbra areas. Note PEF’s RSN membership is CON on the right hemisphere. Data available at: http://balsa.wustl.edu/P2MXl **(b)** Areal responses to each of the three contrasts. A colored X means the area did not survive Bonferroni correction for 360 areas (p<0.05); red n-back, blue stop, green switch). Colors of areal names show RSN membership (color scheme as in a, with the addition of red = DMN). **(c)** Subject overlap map of cortical vertices that were significantly activated in all three executive contrasts for individual subjects (p<0.05 FDR corrected). Black borders surround core MD. Colored borders show RSN membership (CON=blue, DAN=green). Data available at: http://balsa.wustl.edu/7x6l7

Figure 3a shows the RSN membership of all 31 regions. Utilizing the HCP-based 12 network parcellation (Ji et al. 2019), all areas but one belonged to 3 RSNs (FPN, DAN, CON), commonly known in the literature as the three executive networks. We have previously demonstrated core MD forms a strongly interconnected subset of the FPN, splitting the FPN into core and noncore portions (Assem et al. 2020). Only PCV on the medial parietal surface belonged to a different RSN, which Ji et al. (2019) labelled the parietal multimodal network (PMN). Hence, executive activations show unity across multiple “executive” cortical networks, with core MD retaining its strong domain-general properties.

For each core MD, 2020-penumbra and additional-penumbra region, Figure 3b shows results of each individual task contrast (as above, one-sample t-test against zero, p<0.05 Bonferroni corrected for 360 areas). For core MD, the great majority of individual contrasts were positive. The same was true for the subset of 2020-penumbra regions that belonged to DAN or CON. By definition, additional-penumbra regions showed significant contrasts for all tasks in at least one hemisphere, usually both, and notably, the majority of these also belonged to DAN or CON. A further notable result is that, among penumbra regions, those belonging to DAN tended to show greatest activation for switch, while those belonging to CON showed strongest activation for stop.

Next we investigate task overlaps at the finer-grained cortical vertex level. We performed this analysis within-subjects to confirm the existence of overlaps at the single subject level. For each subject, we identified the vertices that were significantly activated across the four runs for each of the three contrasts separately (p<0.05 FDR corrected for cortical vertices). Then we performed a conjunction to identify significant vertices in all three tasks creating a single map per subject. We then summed the maps across subjects to create a probabilistic subject overlap map (Figure 3c). This revealed overlaps in 8 out of the 9 MD patches with little to no overlaps in the temporal MD patch (Figure 3c). Approximately 80% of the vertices that overlapped in 5 or more subjects lay within previously defined and additional MD regions (24.6% core MD, 26.8% 2020-penumbra MD and 29.0% additional-penumbra MD). Vertices with peak overlaps (>70% subject overlap) lay in core MD regions p9-46v, IP2, AVI, and 8BM as well as 2020-penumbra regions SCEF (medial frontal) and AIP (lateral parietal) (Figure 3c).

One intriguing finding is that overlaps lay near borders between core MD and adjacent RSNs in at least 7 locations. In the frontal dorso-medial patch (Figure 3c, bottom 4^th^ column), overlaps traversed the border between 8BM (core MD) and SCEF (CON), an almost identical location to that identified previously using the HCP tasks (Assem et al 2020). In the dorsal lateral frontal patch (Figure 3c, bottom 1^st^ column), overlapping vertices occupy the intersection of i6-8 (core MD), FEF (CON), 6a (DAN). In the two anterior frontal patches (Figure 3c, bottom 1^st^ column), the strongest overlaps lie at the intersection of core MD regions p9-46v and a9-46v with CON regions 46 and 9-46d, respectively. In the insular region (Figure 3c, bottom 2^nd^ column), overlaps lie at the intersection of core MD region AVI with CON region FOP4. In the lateral parietal surface (Figure 3c, bottom 3^rd^ column), the strongest overlaps cross the junction of IP2 (core MD) and AIP (DAN). In the medial parietal surface (Figure 3c), overlaps spanned the junction between POS2 (non-core FPN) and MIP (DAN). Hence, in multiple locations across the cortex, the overlapping vertices lie at the intersection between core MD, DAN and CON. These results suggest that interactions between core MD and adjacent RSNs play a domain-general role in supporting executive functions. We examine these interactions at a finer scale in the following sections. Meanwhile, the conjunction of 3 executive tasks establishes overlapping vertices at the single subject level, especially within and immediately adjacent to core MD regions.

### Diversity of executive functions reflected in canonical RSNs

In the previous section we examined conjunctions across executive tasks. Much previous research, however, emphasizes dissociations between executive functions. Indeed, Figure 1c points to some differences between the tasks and Figure 3b hints that they might be linked to different RSNs. In this section we focus on functional preferences across the three tasks.

To investigate these preferences at a finer grained level, we analyzed data at the vertex level. For each vertex, we compared its activations between the three tasks across subjects (paired t-tests) and assigned the vertex a task label if its activation was significantly stronger than each of the other two tasks (Figure 4a; p<0.05, FDR corrected for cortical vertices and Bonferroni corrected for 3 task comparisons; unthresholded activation group average maps in **Supplementary figure 2** and single subject maps in **Supplementary figure 3**). Figure 4a shows that the most functionally preferred vertices for each contrast surround core MD regions. Some vertices overlapped with our previously-identified penumbra regions (compare Figure 3a), as well as additional regions in a spatial pattern reminiscent of canonical RSNs. For example, comparing task preferences to the HCP-based 12 RSNs (Ji et al 2019; Figure 4b), shows stop>no stop topography (blue) overlaps with areas belonging to the cingulo-opercular network (CON) such as dorsal frontal region 46, inferior frontal area 6r, opercular area FOP5, and angular gyrus region PF. In both hemispheres, 3>1 n-back preferring vertices (red) overlap with the default mode network (DMN), overlapping with dorsal frontal (8Ad), temporo-parietal (PGs), medial parietal (31pd) and medial frontal (s32) areas. On the left hemisphere, switch>no switch vertices (green) overlap with dorsal attention network (DAN) areas in parietal (LIPd) and frontal cortices (6a). Switch also preferentially activates a sliver of left hemisphere vertices along the ventral aspect of frontal core MD regions. The HCP-based parcellation does not include an anterior-ventral frontal portion for DAN. However, fine-grained seed-based examination of an independent HCP resting-state dataset (see **Materials and Methods**) indeed suggests a portion of IFSa, ventral to p9-46v, is strongly connected to DAN (**Supplementary figure 4**). While re-defining fine-grained resting-state cortical networks is beyond the goal of this manuscript, these data nevertheless explain how switch preferences in the ventral portion of the mid-frontal patch are likely related to a fine-grained intrinsic cortical organization.

**Figure 4.**
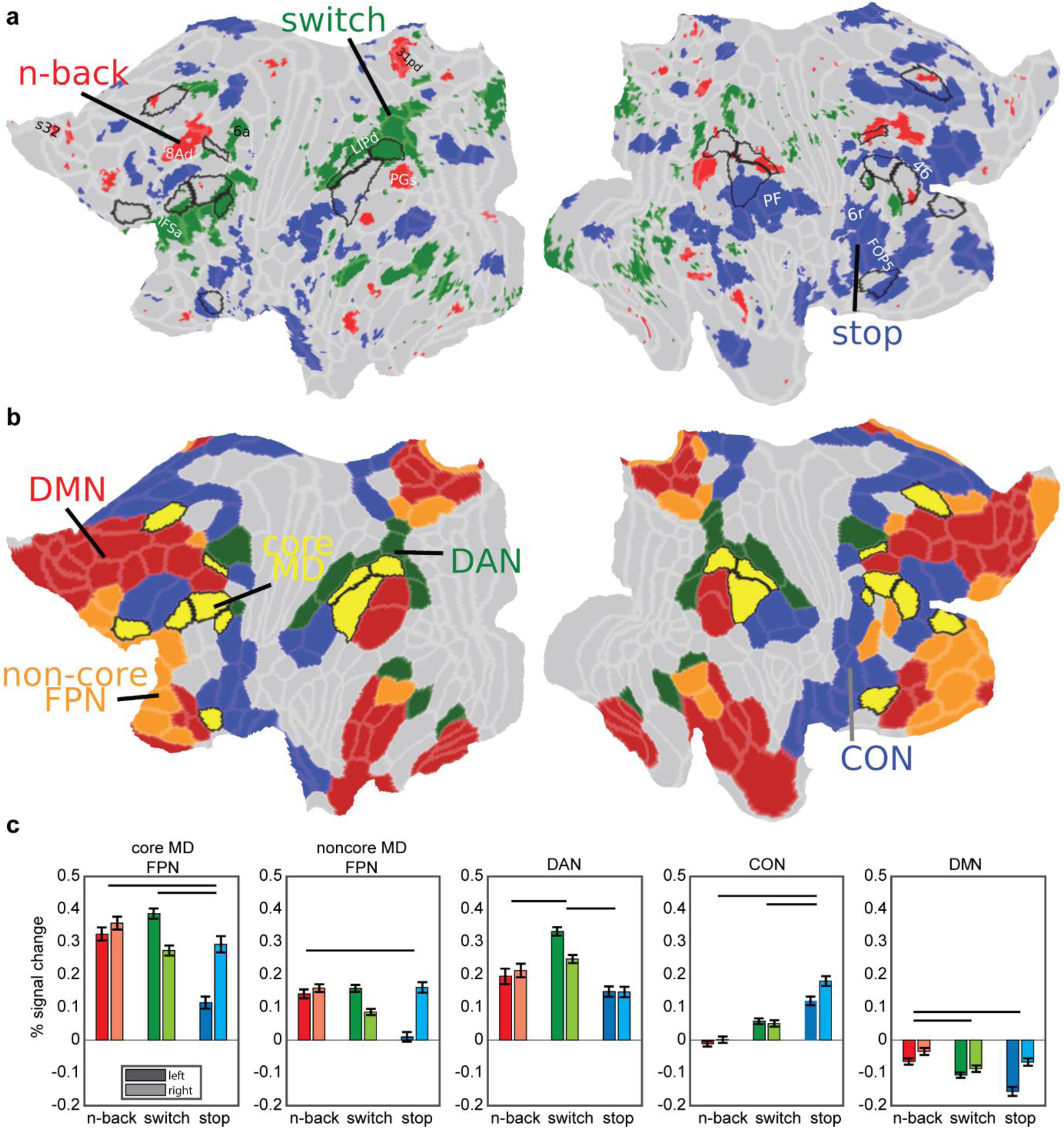
The diversity of executive functions. **(a)** Task functional preferences. Each vertex is colored with the task that significantly activated it more than each of the other two tasks (p<0.05 FDR corrected across vertices and Bonferroni corrected for three tasks; Red: 3>1 n-back, green: switch>no switch, blue: stop>no stop). Core MD areas are surrounded by a black border. **(b)** Canonical RSNs from the HCP based 12 network parcellation by (Ji et al. 2019) (Red: DMN, green: DAN, blue: CON, yellow with black borders: core MD in FPN, orange with grey borders: noncore MD FPN). Note similarity in the topographical organization with each task preference in (a). Data available at: http://balsa.wustl.edu/647Nr **(c)** Task activations for each of the five networks in (b). Error bars are SEMs. Darker colored bars for left hemisphere, lighter colored bars for right hemisphere. Horizontal black lines compare significance between tasks collapsed across hemispheres (p<0.05 Bonferroni corrected for 3 tasks and 5 networks).

To quantify the engagement of RSNs, we used the HCP-based whole-brain definitions of FPN, CON, DAN and DMN to compare activations between the three contrasts (Figure 4c). We split the FPN into core MD and non-core MD portions. Across the five networks, hemispheric asymmetries were pronounced. Collapsing across all networks, 3>1 n-back is right lateralized (paired t-test, t_36_= 8.8, *p*<0.01), stop>no stop is strongly right lateralized (t_36_= 2.9, *p*<0.0001) and switch>no switch is strongly left lateralized (t_36_= 6.6, *p*<0.0001). Within core MD, collapsing across hemispheres, stop>no stop showed the weakest activations (stop vs n-back t_36_=6.5; stop vs switch t_36_=5.5, both *p*s<0.0001), a trend which extended to noncore FPN albeit less prominently (stop vs n-back t_36_=4.8, *p*<0.0001; stop vs switch t_36_=2.3, *p*=0.03). DAN activations were strongest for the switch>no switch compared to 3>1 n-back (t_36_=4.4, *p*=0.0001) and stop>no stop (t_36_=10.0, *p*<0.0001). CON activations were strongest for the stop>no stop contrast compared to n-back (t_36_=9.9, *p*<0.0001) and switch (t_36_=6.9, *p*<0.0001). DMN showed the weakest deactivations during the 3>1 n-back contrast compared to switch>no switch (t_36_=5.2, *p*<0.0001) and stop>no stop (t_36_=4.2, *p*=0.0001). These results show that while each executive contrast strongly activates core MD, each contrast also preferentially activates one or more canonical RSNs, most prominently CON for stop>no stop, DAN for switch>no switch, and reduced DMN deactivation for 3>1 n-back.

Together with the results of the previous section, the unity and diversity of each executive contrast can be conceived as a combination of core MD activations (unity) and adjacent more specialized RSNs (diversity), with different executive demands preferentially recruiting different RSNs. As shown in Figure 3a, however, despite relative specialisations, DAN and CON also show an element of unity, with positive activation across all tasks. In the following section we supplement this RSN-based analysis with examination of fine-grained topographies of executive functions within core MD regions.

### Fine-grained core MD activations are shifted towards task-specific RSNs

An interesting observation from the previous section is that executive preferences, which roughly map out canonical RSNs, surround core MD regions (Figure 4). Here we wondered whether this organization is related to the fine-scaled sub-areal activations within MD regions. We hypothesized that core MD activations for each task would be shifted towards the more specialized RSNs suggesting interaction between relatively domain-specific networks and the closest patches of core MD.

For an initial overview of the fine-scaled preferences, we first selected all vertices that were significantly activated by any of the three contrasts (one-sample t-test against zero; *p*<0.05 FDR corrected for cortical vertices). Then we colored each vertex with the relative strength of its group average activation for each contrast (see **Materials and Methods**). Figure 5a reflects a mosaic of functional preferences, especially within core MD regions. For example, note how the colors form a rapidly shifting gradient within the right p9-46v.

**Figure 5.**
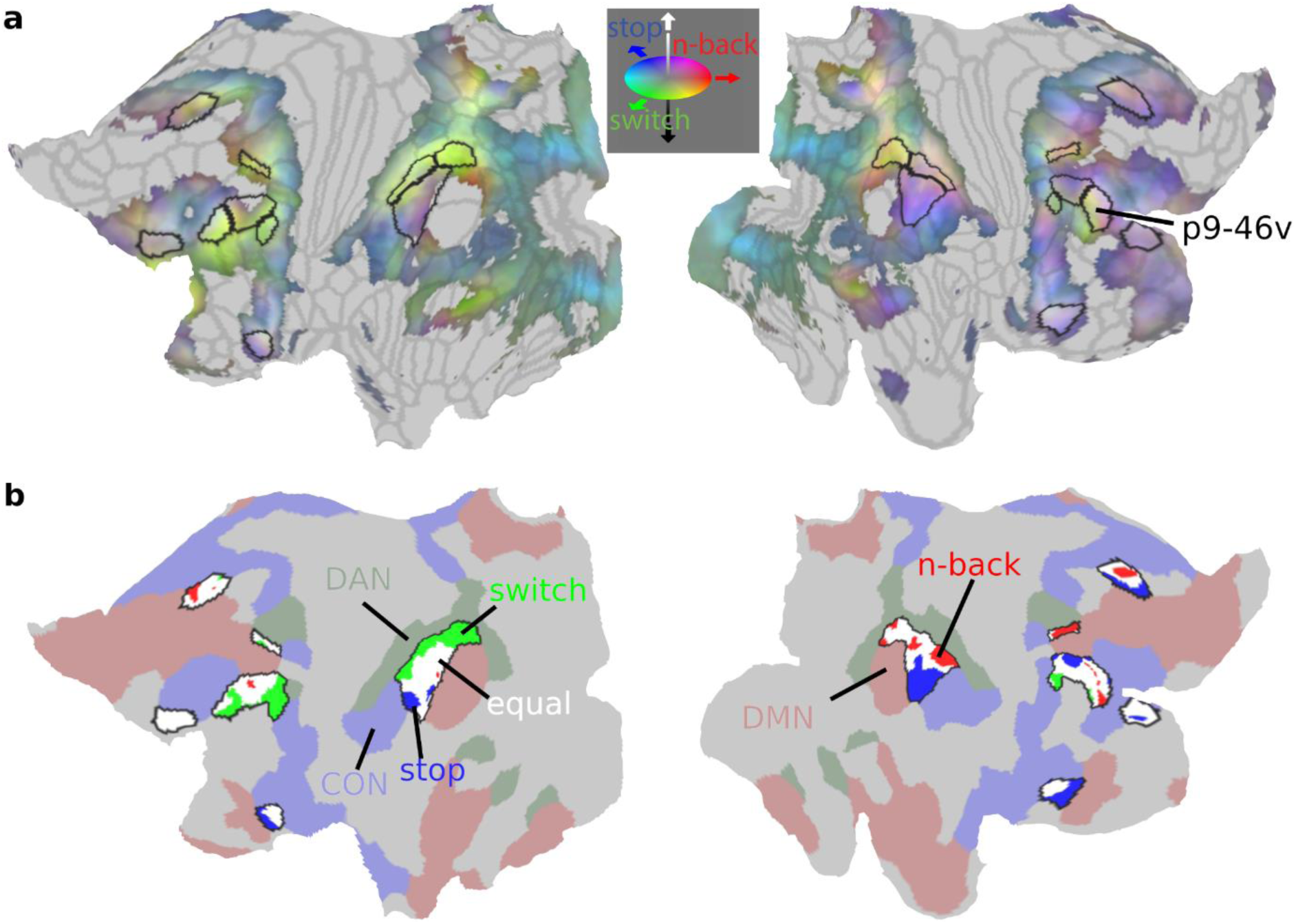
Sub-areal task preferences. **(a)** Cortical projection of the RGB color weighted normalized task profiles. Reddish colors mean stronger n-back activity, bluish colors mean stronger stop contrast activity and greenish colors mean stronger switch contrast activity. Core MD areas are surrounded by black borders. **(b)** Vertex-level statistical comparison of activations within core MD regions. N-back preferring vertices are in red, switch vertices in green and stop vertices are in blue. White vertices denote non-significant statistical differences between tasks (p<0.05 FDR corrected). Surrounding core MD regions (black borders) are canonical RSNs from (Ji et al. 2019) (red: DMN, green: DAN, blue: CON). Data available at: http://balsa.wustl.edu/1p0wB

Next we quantified these preferences within core MD regions using a similar statistical approach to the previous section. We compared activations between the three contrasts at the vertex level and assigned the vertex a task label if its activation was significantly stronger than each of the other two tasks (paired sample t-test, *p*<0.05 FDR corrected for core MD vertices). To facilitate viewing gradients, we combined 3 frontal and parietal regions into two patches (mid-frontal patch: p9-46v, 8C and IFJp; parietal patch: PFm, IP2, IP1). We also visualize the territories for DAN, CON and DMN (Figure 5b).

Figure 5b reveals that many core MD vertices do not have a statistical preference for a specific task. However, among the smaller group of vertices that do show a statistical task preference, they tend to be spatially located closer to their corresponding RSNs. For example, within the parietal patches, switch preferring vertices are closer to DAN within the left hemisphere, and stop vertices closer to CON within the right hemisphere (see **Supplementary figure 5** for single subject examples). These fine-grained trends are less pronounced in frontal patches, likely due to the coarse definitions of canonical RSNs. Altogether, these results demonstrate that fine-grained activations within core MD regions may shift towards different RSNs in a task-specific manner.

### Executive functional preferences are recapitulated by fine-scaled core MD connectivity

The above results reveal fine-scaled activation gradients within core MD regions. Previously, we hypothesized that task preferences likely reflect differences in local MD connectivity with adjacent regions (Assem et al. 2020; Duncan et al. 2020). Thus, here we wondered whether core MD resting-state connectivity is related to activation shifts.

To illustrate, we first focus on the fine-grained preferences of one core MD region in the lateral prefrontal cortex: right p9-46v (Figure 6a). We utilized an independent resting-state data from 210 HCP subjects [the 210V validation sample from (Glasser, Coalson, et al. 2016)]. Each subject underwent 1 hour of resting-state scans (minimally preprocessed, MSMAll aligned, spatial and temporal ICA+FIX cleaned; **see Materials and Methods**) to create a group average vertex to vertex connectivity matrix. We did not use the resting-state data of the current study’s subjects to avoid circularity, as their resting-state data were also used for cortical alignment (MSMAll; see **Materials and Methods**). We then looked at the connectivity patterns of 3 seed vertices, manually placed within right p9-46v (Figure 6a). Seed 1 was placed close to the dorsal p9-46v border with area 46, where stop>no stop preferences peaked (first column in Figure 6a). Intriguingly, the connectivity of seed 1 strongly resembled core MD activations for stop>no stop. In the parietal lobe, for example, connectivity was strongest at the border between areas PFm and PF, with little connectivity to the more posterior IP1. This connectivity-activation similarity extended to other locations throughout the cortex in frontal, parietal and temporal patches (first column in Figure 6a). Next we placed seed 2 roughly in the middle of p9-46v (second column in Figure 6a). Now the connectivity map strongly resembled the n-back contrast activations. On the lateral parietal surface, for example, its peak connectivity lay within intraparietal sulcus area IP2. For seed 3, a vertex near the ventral border of p9-46v (third column in Figure 6a), the connectivity map resembled the switch contrast activations. This qualitative demonstration suggests that fine-grained differences between task activation maps are matched by corresponding fine-grained differences in functional connectivity (Robinson et al. 2014; Tavor et al. 2016).

**Figure 6.**
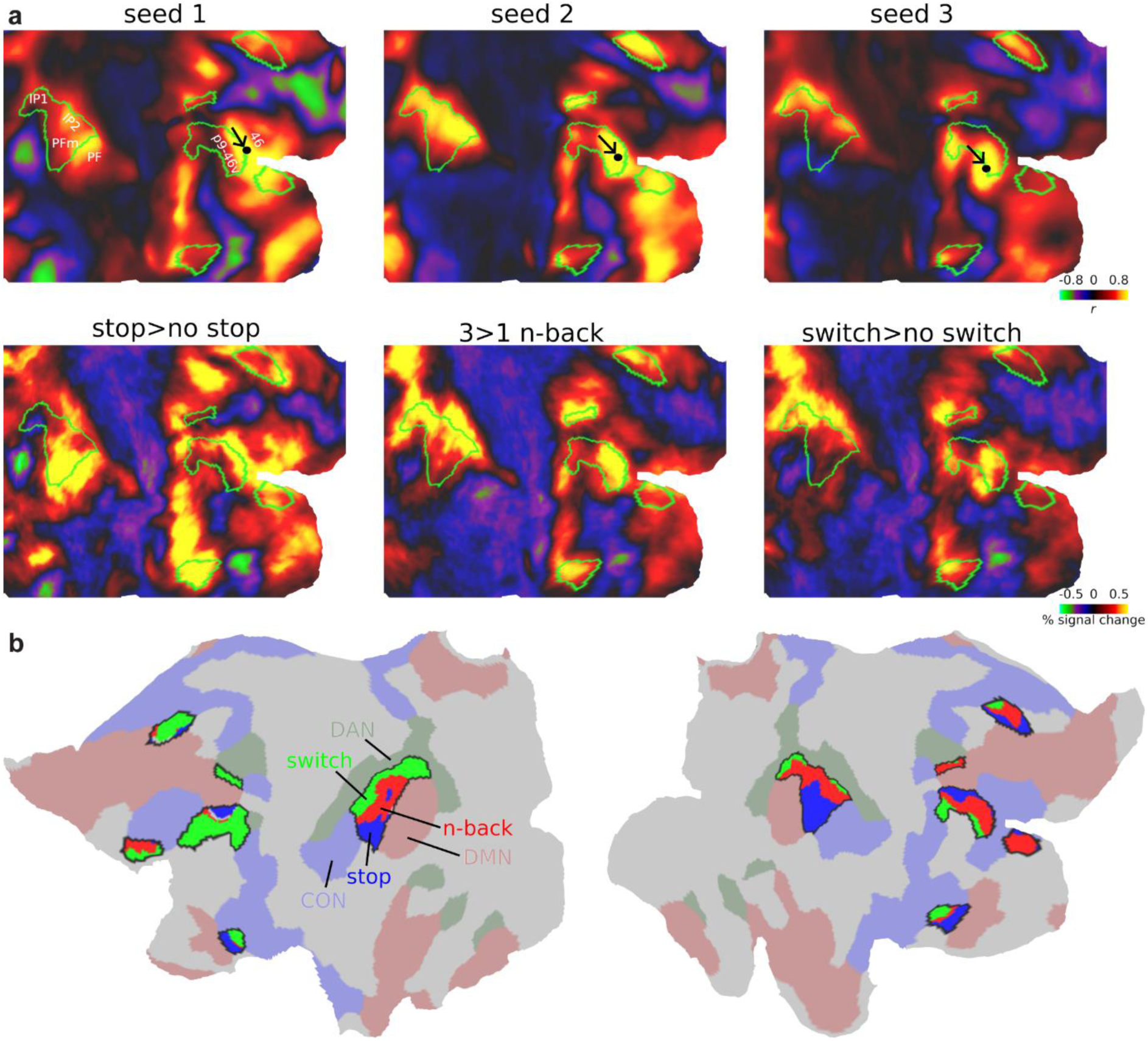
Core MD connectivity gradients. **(a)** Top row: connectivity map of three seeds within right p9-46v. Bottom row: group average activations for each executive contrast. Core MD regions shown with green outlines. Data available at: http://balsa.wustl.edu/5BPVB **(b)** Core MD vertices colored using a winner-take all approach: blue, red, green for vertices where more subjects overlapped for stop, n-back, switch, respectively. Surrounding core MD regions (black borders) are canonical RSNs from (Ji et al. 2019) (red: DMN, green: DAN, blue: CON). Data available at: http://balsa.wustl.edu/n82L9

Next, we quantified the connectivity-activation similarity across core MD regions at the single subject level. For each subject, we correlated (Pearson’s correlation) the connectivity map of each core MD vertex (“seed vertices”) with each of the three activation contrasts. The correlation was performed for seed vertices within core MD regions only. To limit the effect of local connections on driving any correlations, for each seed, we excluded from the correlation all vertices in the seed’s own MMP1.0 region or any adjacent region, along with homologs in the opposite hemisphere. For each subject, we assigned each vertex showing a significant correlation greater than 0.2 with any of the task maps (FDR *p*<0.05) the label of the task it numerically most strongly correlated with. Then we created a subject probabilistic map for each task. To facilitate viewing the connectivity gradients within core MD regions, we used a winner-take all approach to assign each core MD vertex the label of the task with the greatest subject overlap (Figure 6b). The results reveal fine-scaled systematic shifts in connectivity gradients. For example, replicating our manual seed demonstration, a dorsal to ventral task gradient (stop to n-back to switch) exists within the right mid-frontal MD patch. The gradient is reversed to a ventral to dorsal direction in parietal MD regions. In the insular patch, it becomes rostral to caudal. Note how seed preferences often follow the canonical RSNs; switch seeds closer to DAN (green), stop seeds closer to CON (blue) and n-back seeds closer to DMN (red).

Collectively, these results demonstrate fine-scaled topographies of each executive demand can be predicted by fine-scaled core MD connectivity. Hence, co-activated vertices within core MD form functionally connected networks. These findings further support the notion that each executive demand recruits a task-specific network which in turn interacts most strongly with its immediately adjacent core MD territories.

### Activations peak at borders between core MD and adjacent RSNs

Within the fine-scale patterns that we have shown, borders seem especially important, including borders shared between core MD and other RSNs. For example (Figure 6), stop activations peak at the border between p9-46v (core MD) and area 46 (part of CON). Here we wondered whether there is a general pattern where peak activations fall on the borders between core MD and task-relevant adjacent networks.

First, we wanted to map out, at the single subject level, where the strongest activations lie for each contrast. For each subject and task separately, we selected the top 5% activated vertices across the whole cortex, binarised the map and summed it across subjects to create a probabilistic subject overlap map for each task. Vertices that overlap in more than 50% of subjects are shown Figure 7a. The strongest overlaps are concentrated around p9-46v, the parietal patch, and medial parietal patch. Stop>no stop vertices were closest to borders between core MD and CON, switch and n-back vertices were closest to borders between core MD and DAN.

**Figure 7.**
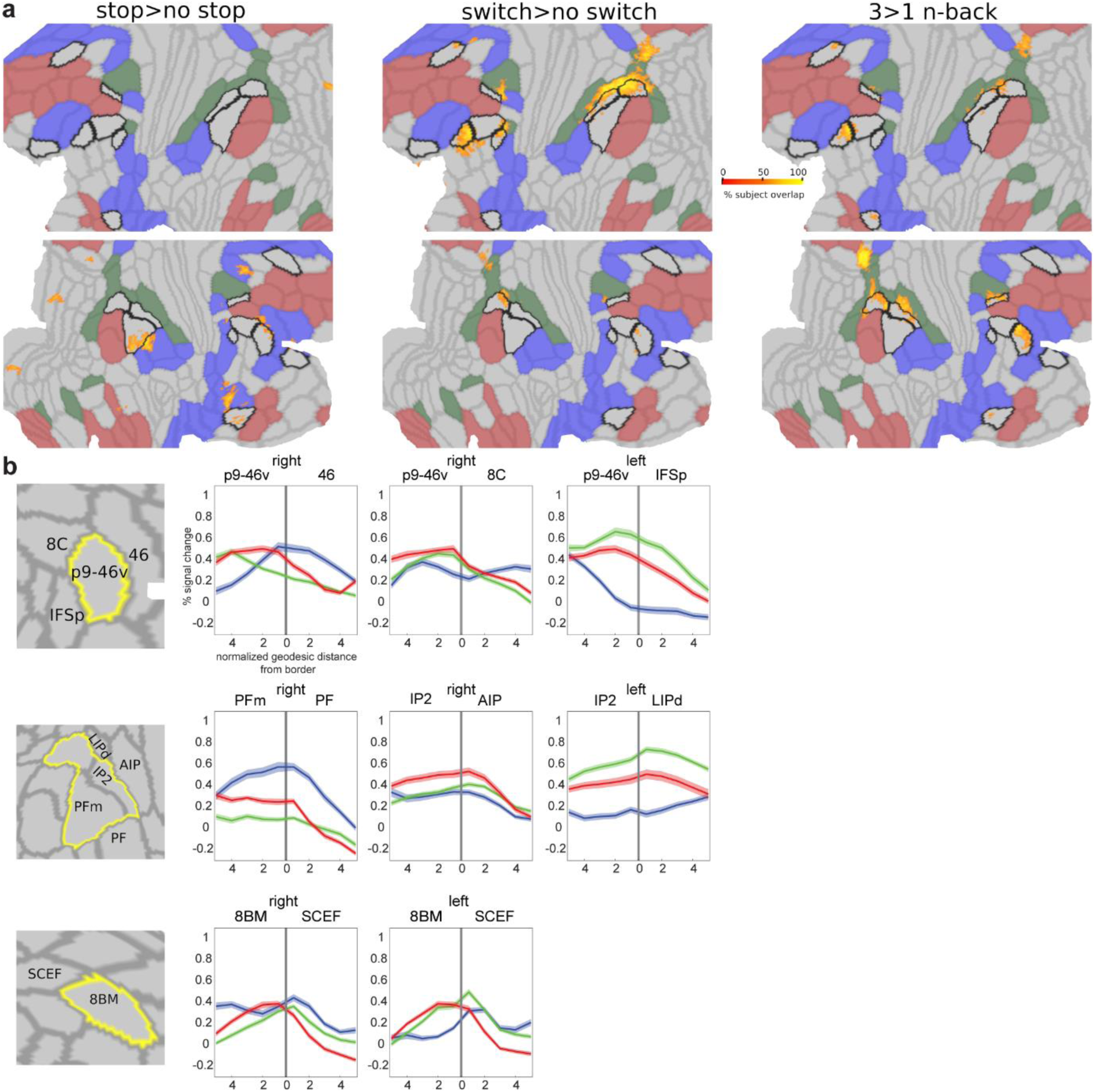
Peak activations at region borders. **(a)** Subject overlap map of top 5% activated voxels for each contrast. Core MD borders are colored in black and the remaining MMP1.0 areas borders are in light grey. RSNs are colored as follows: DMN is red, DAN is green, and CON is blue. Data available at: http://balsa.wustl.edu/gmNwX **(b)** First column represents a close up of cortical areas of interest (top lateral prefrontal, middle: lateral parietal, bottom: medial frontal). Remaining columns display example activation profiles near areal borders, highlighting the hemisphere –right or left-with the strongest pattern. N-back activations are in red, stop in blue and switch in green. Shaded areas represent SEMs. The location of the border is marked by a vertical grey line at the zero point of the x-axis.

To get a better picture of activations at border zones, we analyze borders between p9-46v and its surrounding three regions: 46, 8C, IFSp (Figure 7b**; top)**. For each pair of regions, we divided the vertices within each area into 10 equal segments based on their geodesic distance from their shared border (see **Materials and Methods**) and statistically compared their activations along the 10 segments across subjects (one way repeated measures ANOVA). A significant segment x task interaction for all pairs (all F_(30,1080)_>128, *p*<0.0001) show activations peaking at borders that were strongest for their corresponding contrasts (stop strongest at right p9-46v/46, switch at left p9-46v/IFSp, n-back at right p9-46v/8C).

We also examined activation peaks at the borders of the parietal core MD patch (Figure 7b**; middle**). The stop contrast peaks near the border of PFm and PF (CON) on the right hemisphere, n-back peaks near the border of IP2 and AIP (DAN) on the right hemisphere and switch peaks near IP2’s border with LIPd on the left hemisphere (all segment x task interactions F_(30,1080)_>89, *p*<0.0001). This follows the tight intrinsic functional relationship between mid-frontal and parietal patches we demonstrated in the previous section (Figure 6).

On the medial frontal surface, our previous work (Assem et al. 2020) noted a striking consistency across all tasks, with peak activation at the intersection of core MD area 8BM with SCEF (CON) (Figure 1). Probing this intersection in the current data (Figure 7b**; bottom**) indeed confirmed that activations for all three tasks peak near or at the 8BM/SCEF border bilaterally (all segment x task interactions F_(30,1080)_>78, *p*<0.0001).

As the above results rely on group-defined HCP_MMP1.0 borders, we sought to alleviate concerns that border activations might simply reflect an artefact due to individual differences in border locations or inherent MRI signal smoothness. To test this, we created a simulated dataset utilizing the subject-specific cortical parcellations of a randomly selected 37 HCP subjects from the 449 subjects dataset (see **Materials and Methods**). For each subject and for each area, we assigned all its vertices the activation value of the corresponding area and subject in the real data. This simulates the condition that activations are homogenous throughout the area. We created three datasets for our three contrasts. We then applied several degrees (4, 12, 20 mm FWHM) of surface smoothing to simulate the inherent blurriness of fMRI data at varying levels. Then we repeated the same analysis above, selecting the top 5% activated vertices for each subject and creating an overlap map across subjects. While the simulated maps showed activations in the same zones, none of the simulated maps replicated the sharp border activations. For example, comparing the simulated data (12 mm) with the real data shows much broader activations, with little focus on regional borders (**Supplementary figure 6**). With 4 and 20 mm smoothing, results even less resembled the actual data.

These spatially precise results suggest that areal borders play a critical role in communication between MD and adjacent networks. Likely, it is at these borders that information is most intensively integrated between networks.

### Overlap and divergence in subcortical and cerebellar MD regions

Finally we wondered whether executive overlaps and preferences extend to subcortical and cerebellar MD regions. We focus on the caudate, thalamus and cerebellum as those were the most prominent extracerebral structures that showed MD properties in our previous study (Assem et al. 2020). The three structures were segmented for each individual separately as part of FreeSurfer’s standard segmentation of 19 subcortical/cerebellar structures (see **Materials and Methods**). This avoids mixing of signals from the white matter or between different structures.

To identify overlaps at the voxel level, for each structure, we identified significantly activated voxels for each contrast (one sample t-test, FDR corrected for each structure separately, *p*<0.05, Bonferroni corrected for 3 structures) and then identified the conjunction of significant voxels across the three contrasts (Figure 8a). For analysis transparency, we report that the initial results using the unsmoothed data (i.e. no additional smoothing over the standard 2 mm at the preprocessing stage) for the caudate and thalamus were patchy, likely due to the relatively low signal to noise ratio (SNR) of high spatial and temporal resolution fMRI for the subcortex at 3T because these regions are far from the head coil. Thus here, for caudate and thalamus, we report the results after applying a 4 mm FWHM smoothing kernel limited within the major subcortical structures. This revealed a cluster of voxels in caudate head, extending to its body bilaterally, overlapping with caudate MD regions identified by Assem et al. (2020) (Figure 8b, yellow). We were also able to identify a thalamic cluster that significantly overlapped with the putative MD thalamic region from Assem et al. (2020). For the cerebellar analysis we did not apply any additional smoothing, as the cerebellum is generally closer to the head coil. The analysis revealed clusters located mainly in cruses I and II, again mainly overlapping with MD cerebellar regions. Note however, that two of the previously-defined MD patches occupying the medial and lateral portions of the right crus I are missing in the current data. Investigating each contrast separately (**Supplementary Figure 7**) revealed that this is due to deactivations in this region for the stop>no stop contrast. These results largely replicate our prior findings of MD regions in subcortex and cerebellum.

**Figure 8.**
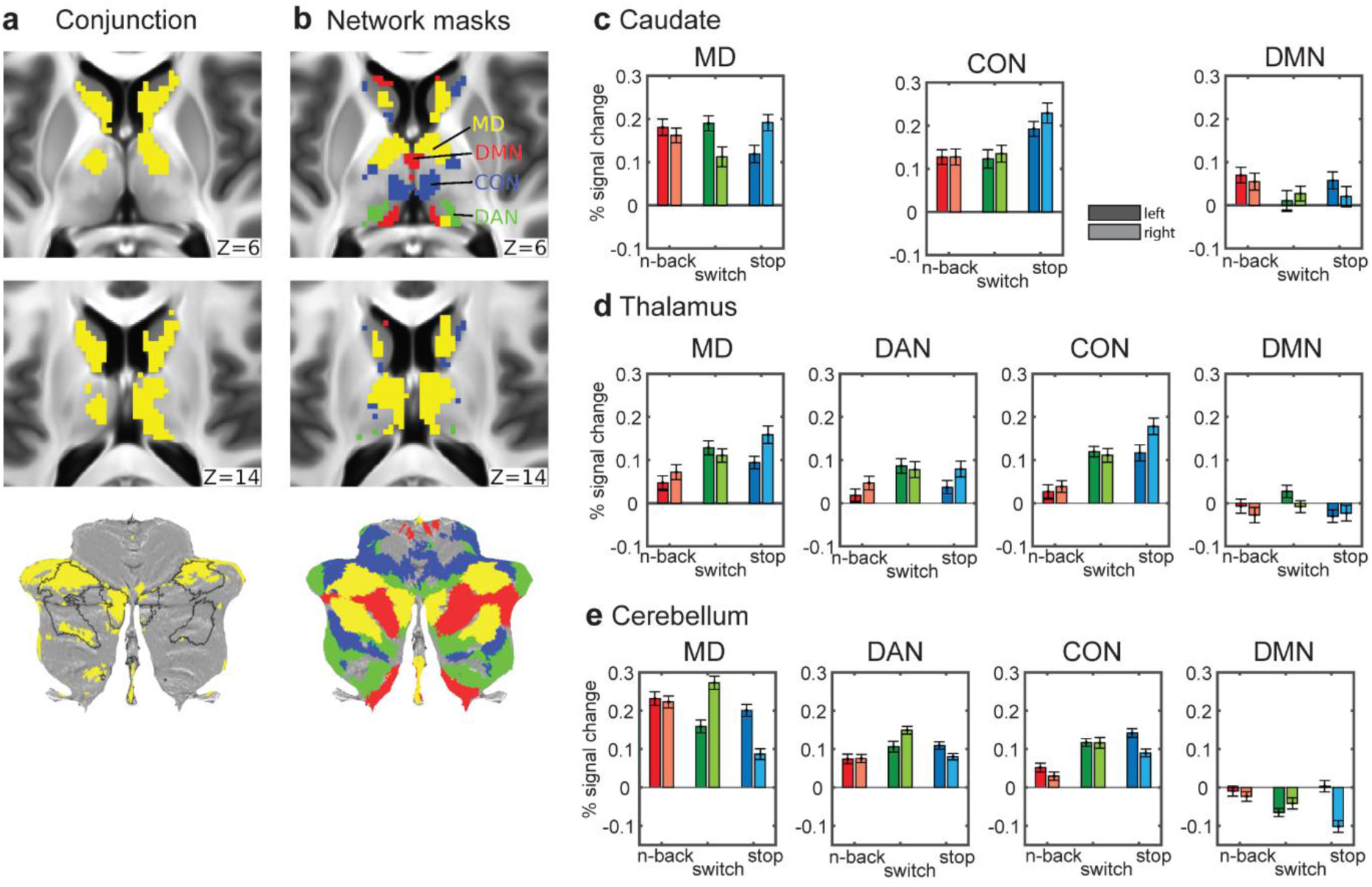
Conjunctions and task preferences in subcortex and cerebellum. **(a)** Subcortical axial slices and a cerebellar flat map showing surviving voxels in caudate, thalamus and cerebellum for the conjunction of significantly activated voxels for each of the three executive contrasts (p<0.05 FDR corrected for each structure separately). **(b)** Voxels belonging to each network (yellow=MD, green=DAN, blue=CON, red=DMN). MD caudate, thalamic and cerebellar voxels are from (Assem et al. 2020). The other three RSN definitions are from (Ji et al. 2019) [see **Materials and Methods** for more details]. Data available at: http://balsa.wustl.edu/Bgp6w **(c), (d), (e)** Bar plots of activations across subjects for each hemisphere, network and structure. Darker colored bars are for left hemisphere and lighter colored ones for the right hemisphere. Error bars are SEMs.

We then wondered whether subcortical and cerebellar activations exhibit functional preferences within MD and adjacent RSNs. For this analysis we used unsmoothed data for all structures. We utilized our subcortical and cerebellar MD definitions (see **Materials and Methods**) and the DAN, CON and DMN subcortical and cerebellar network definitions from the same network parcellation we used for the cortical analysis (Ji et al. 2019). Any overlapping voxels between MD and RSNs were assigned to MD areas. We averaged the voxel activations for each network separately to give one parameter estimate per network x lateralization x structure x subject. To assess statistical significance, we applied a separate ANOVA for each structure with three factors: 3 tasks x 4 networks x 2 hemispheres. The caudate only had 3 networks as it did not have any voxels identified as DAN. All post hoc t-tests were evaluated at *p*<0.05 Tukey-Kramer corrected.

For the caudate (Figure 8c), the task x hemisphere x network interaction was significant (F_(8,288)_=14.0, *p*<0.0001). Post hoc tests confirmed that within the MD area, switch>no switch was left lateralized, stop>no stop was right lateralized and 3>1 n-back showed no hemispheric preferences. Within CON, stop>no stop was significantly right lateralized and most strongly activated. Neither MD nor DMN regions of caudate showed overall task preferences.

For the thalamus (Figure 8d), the interaction task x hemisphere was significant (F_(24,864)_=38.7, *p*<0.0001). Across networks, stop was right lateralized while switch was left lateralized.

For the cerebellum (Figure 8e), we observed a significant interaction between task x hemisphere x network (F_(24,864)_=92.9). Across all networks, post hoc analysis showed left lateralized activations for stop>no stop, right lateralized for switch>no switch but no significant hemispheric preference for 3>1 n-back. These trends were especially strong in the MD area, which also showed strongest activations overall. Note the flipped hemispheric preferences in the cerebellum due to the decussation of fibers across the midline in the brainstem.

All in all, the three executive tasks do share common activations that overlap with MD regions in the caudate, thalamus and cruses I and II in the cerebellum. Within MD, hemispheric differences were more prominent than task preferences. Outside of MD, DAN was preferentially activated by switch in the thalamus and cerebellum, while CON was predominantly activated by stop>no stop. Overall, these results parallel cerebral cortical findings, confirming that task overlaps and dissociations extend to subcortical and cerebellar components of brain networks.

## Discussion

Decades of spatially coarse brain imaging results have left many open questions on the link of executive functions to the functional organization of association cortices. Our results using the high spatial resolution of HCP multimodal MRI approaches provide a novel framework supporting the unity and diversity model of executive functions and bridging it with detailed functional anatomy of the human brain. Activations of three distinct executive functions showed overlapping activations (at the single subject and single vertex/voxel level) within cortical, subcortical and cerebellar domain-general MD regions. Surrounding this unity, each executive demand shows unique functional preferences within MD regions that extend to nearby canonical RSNs. Linking this unity and diversity are strong activations at the intersection of core MD and adjacent task-specific RSNs. We discovered these activations often peaked at network borders defined using multimodal MRI criteria, suggesting a likely substrate for integration between networks. Our novel framework suggests domain-specific areas recruit adjacent MD areas from different spatial locations on the cortical sheet to generate executive functions, likely far more diverse than the three studied here. We elaborate on these points below.

### MD patches: A consistent topology with task-specific shifts

Using the precise HCP imaging approach, we have previously delineated 9 coarse cortical patches (Figure 1) co-activated by 3 cognitively demanding tasks (Assem et al. 2020). In this study we show that each of the three executive tasks strikingly co-activate roughly the same 9 MD territories (Figure 1). An exception was activity in the temporal patch, in which stop activations were more anteriorly-dorsally shifted than the other two contrasts. More generally, within the MD patches, each task showed detailed topological shifts. Our results showed that many of these shifts were unique for each task and varied in a systematic pattern across the cortex linked to the underlying fine-grained functional connectivity.

### Executive unity at the intersection of MD core with adjacent networks

We previously linked MD patches to a set of 10 core MD regions, which roughly outline the central portions of 7 out of the 9 patches. The conjunction of the three executive contrasts replicated 9 out of the 10 core MD regions, with the exception of posterior intra-parietal region IP1, confirming the strong domain-generality of MD core (Figure 3b). Surrounding MD core, the conjunction also identified a number of penumbra areas belonging to three RSNs: CON, DAN and non-core FPN (Figure 3b).

At the finer grained vertex level, the most consistent conjunctions fell at the intersection of MD core with CON and DAN in 6 locations and at a 7^th^ location on the medial parietal patch between non-core FPN (POS2) and DAN. The location of these overlaps is reminiscent of an earlier study which argued for defining cortical integrative hubs at the points of intersection of multiple networks (Power et al. 2013). A follow up study found that selective damage to these hubs is associated with large deficits in executive abilities (Warren et al. 2014). This suggests that our identified overlaps between these 3 networks are critical for supporting executive functions, at least the three components of executive function tested here.

The longstanding debate on the existence of common executive activations in fMRI studies often attributes task overlaps to the merging of distinct, fine-grained networks due to low resolution (Braga et al. 2019). Our work challenges this with some of the highest spatial resolution in the field, using 2 mm voxel resolution and 2 mm FWHM surface-based smoothing. Moreover, overlaps in executive-like tasks have been observed at the single-neuron level in putatively homologous MD areas in non-human primates (Panichello and Buschman 2021). These findings argue that the spatial commonalities in task activations are not solely due to low resolution, suggesting shared neural resources recruited during executive tasks.

### Executive diversity reflects distinct interactions between domain-specific networks and core MD

Many previous fMRI studies focused on dissociations between executive functions, suggesting they are supported by functionally distinct territories of the association cortices (Wager et al. 2005; Dosenbach et al. 2006; Dodds et al. 2011; Hampshire et al. 2012; Eisenreich et al. 2017). Such fractionated conceptions offer limited understanding for how executive processes are integrated and coordinated across the brain.

Instead of functional fractionations, our anatomically resolved results reveal a different picture of a common MD territory, defined by multimodal neurobiological criteria, which can be recruited from different spatial locations on the cortical sheet according to different task requirements. Here we summarize how each contrast’s activation demonstrated this pattern in a spatially unique manner (Figures 4-7).

The stop>no stop activations were strongest in the right hemisphere, a well-documented hemispheric bias (Corbetta and Shulman 2002; Apšvalka et al. 2022) that might be explained by richer arousal-related neuromodulator projections to the right hemisphere (Arvin et al. 1978). Many regions of CON showed stronger activation for stop than the other two tasks (Figure 4a**, b)**, and within core MD regions, activations were often shifted towards adjacent CON regions, especially in parietal cortex (Figure 5). In some cases activations peaked at the boundary between adjacent core MD and CON regions (Figure 7). Previous studies implicate at least four cortical nodes in stop signal or similar paradigms: dorsal frontal regions, inferior frontal junction, temporal-parietal junction and dorsal anterior cingulate (Swann et al. 2012; Aron et al. 2014; Yeo et al. 2015; Suda et al. 2020; Isherwood et al. 2021; Sebastian et al. 2021). While an accurate comparison with our results is not possible, we show that these coarse nodes likely lie at the intersection between core MD and CON. We suggest that interaction between these two networks underlies the attentional re-orienting or “braking” processes involved in stop signal and similar paradigms.

The n-back contrast was slightly right lateralized, showing strong activation throughout core MD (Figures 4**, 5**) with strongest activations shifted mainly towards core MD-coreMD, coreMD-DAN or coreMD-DMN borders (Figure 7). N-back also showed the least deactivation of DMN, with the strongest effects in inferior parietal, medial parietal and dorsal frontal portions, a canonical DMN fractionation usually implicated in memory recall and spatial imagery studies (Figure 4 **& Supplementary Figure 2**) (Andrews-Hanna et al. 2010; Wen et al. 2020; Shao et al. 2023). A recent study also demonstrated co-engagement of intermediate nodes between FPN and DMN in a 1-back>0-back task (Murphy et al. 2018). Murphy et al suggested the interaction between FPN and DMN plays a role in recalling detailed information from the immediate past. We speculate that the engagement of the intersection between parietal DMN (e.g. PGs, part of 2020-penumbra) and core MD patches reflects engagement of the episodic recall network (Rugg and Vilberg 2013) though this needs confirmation through a recall-focused paradigm investigated in a spatially precise approach similar to the current study.

Switch>no switch activations were left lateralized, with strongest activations in core MD and DAN (Figure 4c), in line with previous results highlighting dorsal fronto-parietal activations (Braver et al. 2003; Crone et al. 2006; Tsumura et al. 2021). Intriguingly, the switch contrast also showed strong activations in a band of lateral frontal regions ventral to core MD regions (Figures 1**, 4**). In a parallel analysis, we showed that seeds in these frontal patches show strong connectivity to the dorsal components of DAN, suggesting a ventral frontal component to DAN. The functional role of this ventral component remains unknown. On one hand, this patch of cortex is functionally heterogeneous, with sensory-biased responses (Michalka et al. 2015; Assem et al. 2022) and language responses (Ji et al. 2019). On the other hand, it is consistently engaged when comparing simple cognitive tasks to rest (Assem et al. 2020). The left hemisphere bias and preference of ventral frontal regions may indicate a role for the phonological loop of working memory (inner speech) in managing task switching and wider cognitive tasks.

Thus, the diversity of executive task activations paints a picture of specialized recruitment of core MD regions from adjacent more domain-specialized networks. This adds to accumulating evidence of other domain-specific language and sensory biased regions that also lie adjacent to MD regions (Fedorenko et al. 2012; Assem et al. 2022). These domain-specific regions likely form communication bridges with core MD, feeding in and out task-relevant information to support brain-wide cognitive integration (Duncan et al. 2020). In line with this hypothesis, our functional connectivity analysis demonstrates activation shifts within core MD are mirrored by fine-grained shifts in whole-brain connectivity. These results echo previous demonstrations of pre-frontal activations being constrained by its intrinsic functional architecture (Tavor et al. 2016; Waskom and Wagner 2017).

In this study we looked at three traditional elements of the unity-diversity view. An intriguing question concerns how the results might change with a more diverse set of tasks? Our conception of MD core as a domain-general territory commonly recruited from different spatial directions suggests that diversity is likely far greater than identified here. Activations for diverse cognitive demands are likely to intersect at different junctions between core MD and other RSNs.

### Core MD borders are critical for executive functions

One of the most striking findings in this study is that the three executive contrasts showed peak activations overlapping with borders between core MD and adjacent RSNs (Figure 7). The implicated borders largely follow the RSNs most relevant to the contrast. Stop>no stop peak activations mostly overlapped with core MD-CON borders, while core MD-DAN borders were mostly crossed by switch>no switch peak activations. 3>1 n-back had a mixed preference towards core MD-core MD, core MD-DAN and core MD-DMN borders.

That said, there were also borders where the three contrasts peaked together. Most prominently this occurred in the dorsomedial frontal patch at the border between 8BM (core MD) and SCEF (penumbra). We have also highlighted peaks at this border in our previous study that employed a different set of task contrasts (n-back, reasoning, math>story) (Assem et al. 2020). This striking consistency suggests a precise anatomical correlate for a domain-general process, and is most likely linked to response selection activations in the dorsal anterior cingulate cortex (Si et al. 2021; Seghezzi and Haggard 2022).

These findings are the most detailed in a growing body of evidence of activations lying at network borders (Nee 2021). For example, a recent study highlighted that FPN borders with other networks are the most predictive of individual differences in executive abilities (Reineberg et al. 2022). Why would activations peak at borders? The HCP MMP1.0 areal borders were defined using robust multimodal architectural and functional criteria including cortical curvature, myelin content, functional connectivity, and task activations as well as careful cross examination with previous cyto-architectural studies (Glasser, Coalson, et al. 2016). This suggests a neurobiologically relevant function for borders. Just as areal borders between visual regions reflect shifts in topographic organization of the visual field, borders in association cortices might reflect shifts across the topographic organization of higher cognitive functions. Here we suggest that border activations might reflect the most intensive locations for information exchange between MD and domain-specific regions. More detailed examination of anatomical connections (e.g. using tract-tracing in non-human primates) and neural dynamics (e.g. using invasive electrophysiological recordings) of border zones could further clarify their role in information integration.

### The role of the MD subcortical and cerebellar regions in executive control

Task overlaps and divergences also extended to subcortical and cerebellar regions. The three executive contrasts overlapped with MD regions in the head of caudate, the anterior and medial thalamus, and cerebellar cruses I and II (Figure 8). Our previous study (Assem et al. 2020) failed to find overlapping thalamic activations across diverse tasks, likely due to the low SNR of high spatial and temporal resolution fMRI in deeper brain regions (Assem et al. 2020), but such overlap is clearly visible in the current data. Interestingly, RSN functional preferences also extended into the subcortex and cerebellum, with hemispheric biases and RSN preferences largely matching those of cerebral cortex. These results strengthen the view that the MD system is brain-wide and tightly integrated. To examine finer-grained shifts and link activations with specific subcortical/thalamic nuclei, higher resolution and higher SNR studies with 7T are needed.

### Recruiting a common MD territory to create distinct executive functions

Our results bring a fresh, anatomically precise perspective on brain systems underlying unity and diversity of executive functions. This perspective deviates from the broad differentiations typically observed in lesion studies and functional imaging investigations. We suggest that many cognitive demands, including traditional executive demands, recruit activity in a characteristic territory of 9 cortical patches, with associated subcortical and cerebellar activity. Different demands, however, shift the detailed pattern of activity within these patches, often towards adjacent, more specialized RSNs recruited by individual tasks. With MD territory at the center of multiple domain-specific networks, it is well placed to integrate the components of a cognitive operation to generate distinct executive processes.

## Acknowledgments

J.D was funded by a Medical Research Council grant (MC_UU_00030/7); M.A was funded by Cambridge Commonwealth European and International Trust (Yousef Jameel scholarship); S.S. was funded by Gates Cambridge Trust (Cambridge, UK); M.F.G was funded by R01MH060974.

## Data availability

Data used for generating each of the imaging-based figures are available on the BALSA database (https://balsa.wustl.edu/study/0qk6K). Selecting a URL at the end of each figure will link to a BALSA page that allows downloading of a scene file plus associated data files; opening the scene file in Connectome Workbench will recapitulate the exact configuration of data and annotations as displayed in the figure.

---

**Supplementary Figure 1.**
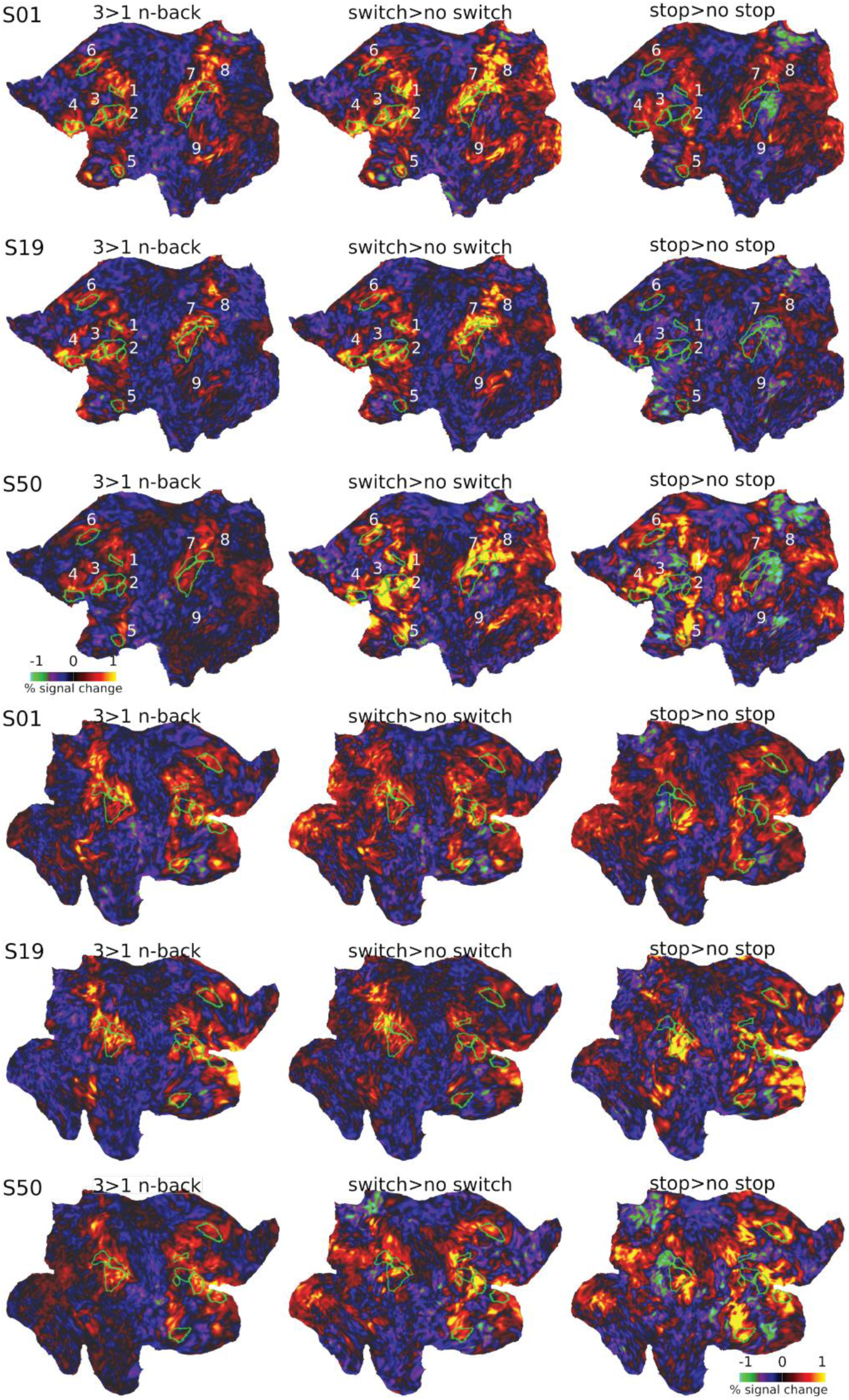
Activation maps for each task for three example subjects. Top three rows are left hemisphere. Bottom three rows are the right hemisphere. Core MD borders are outlined in green. Data available at: http://balsa.wustl.edu/Nkm05

**Supplementary Figure 2.**
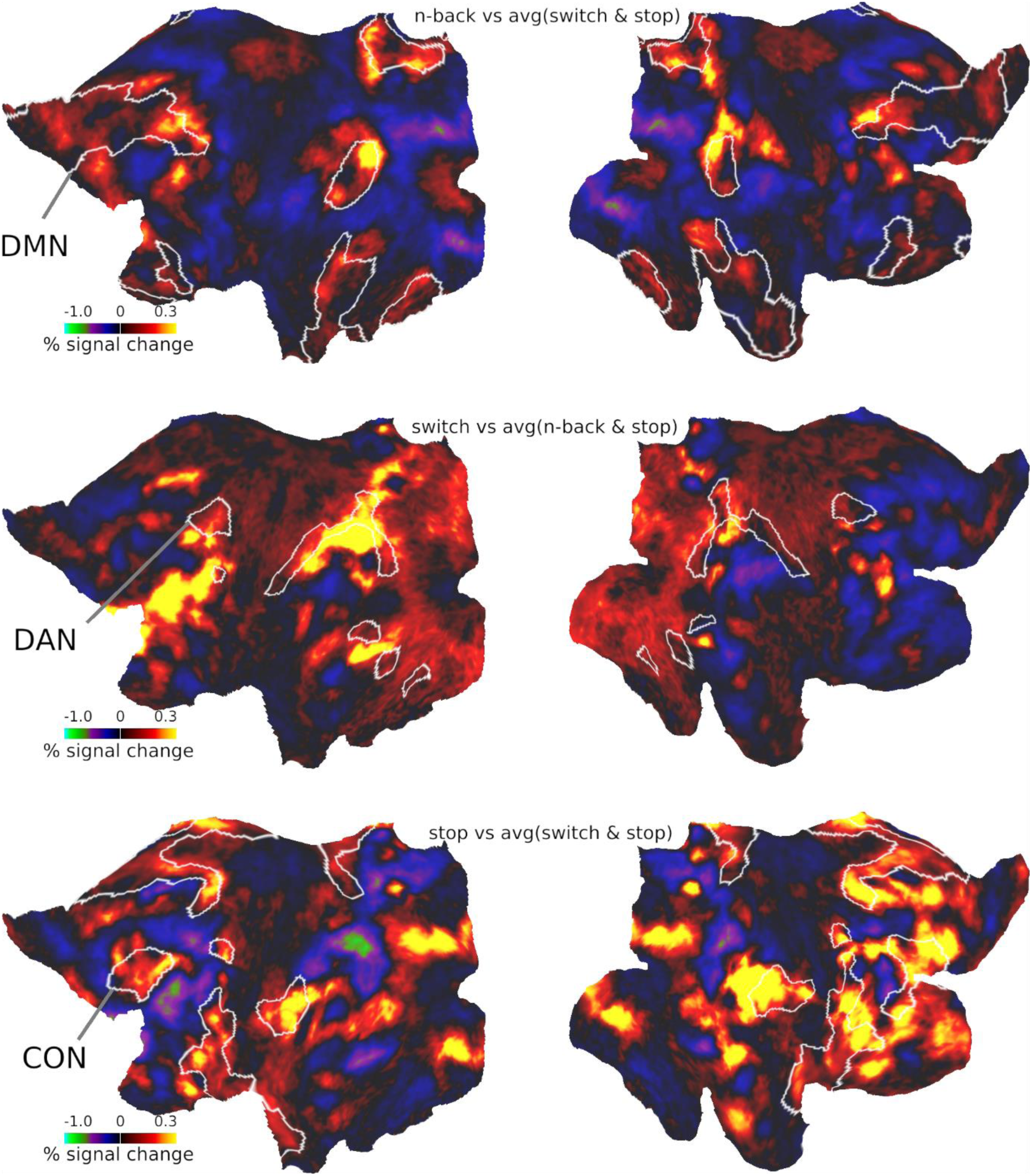
Group average activations of each executive task minus the average of the two other tasks. RSN borders are outlined in white. Data available at: http://balsa.wustl.edu/l7kN9

**Supplementary Figure 3.**
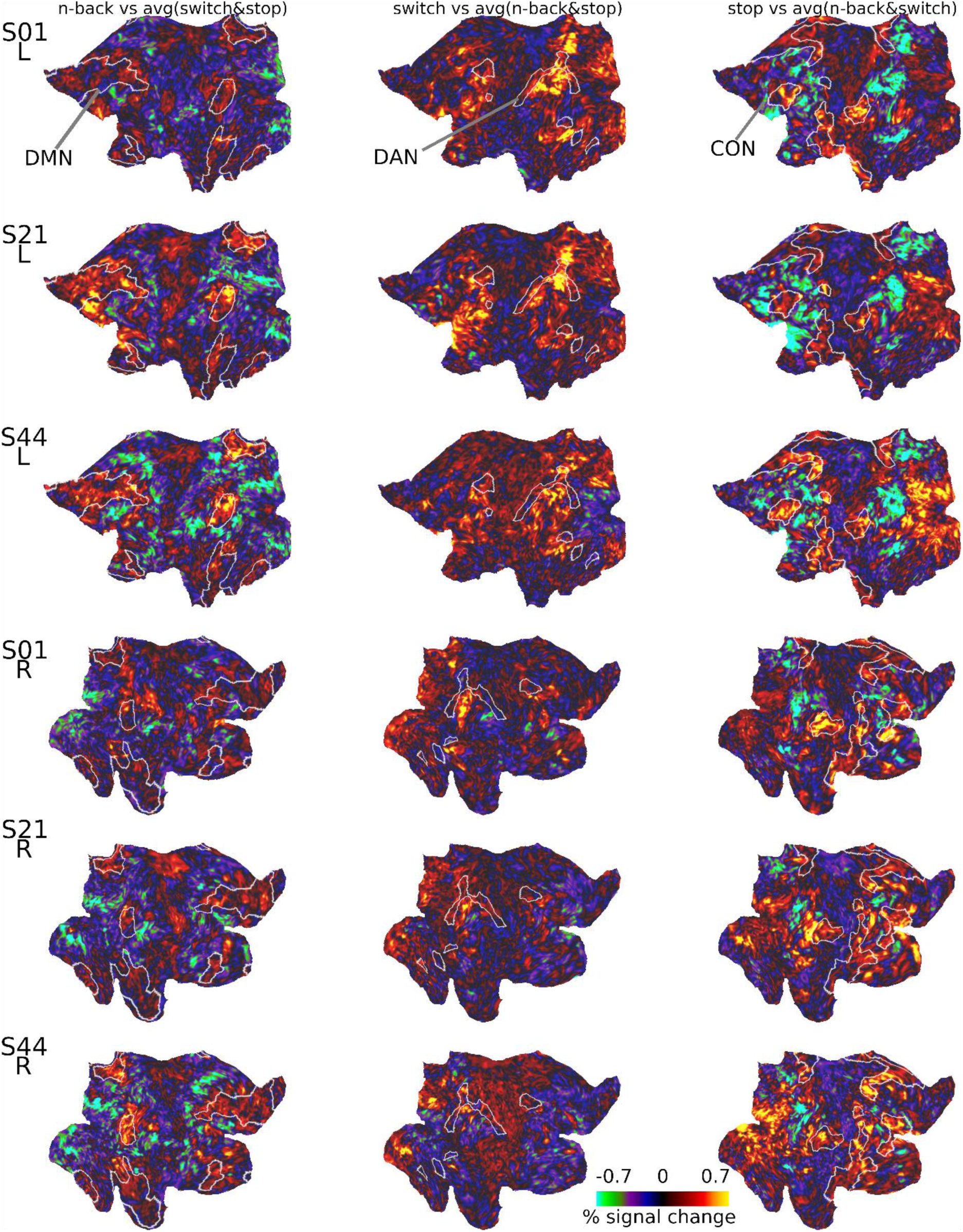
Three example subject activations of each executive task minus the average of the two other tasks. RSN borders are outlined in white. Data available at: http://balsa.wustl.edu/G5P2G

**Supplementary Figure 4.**
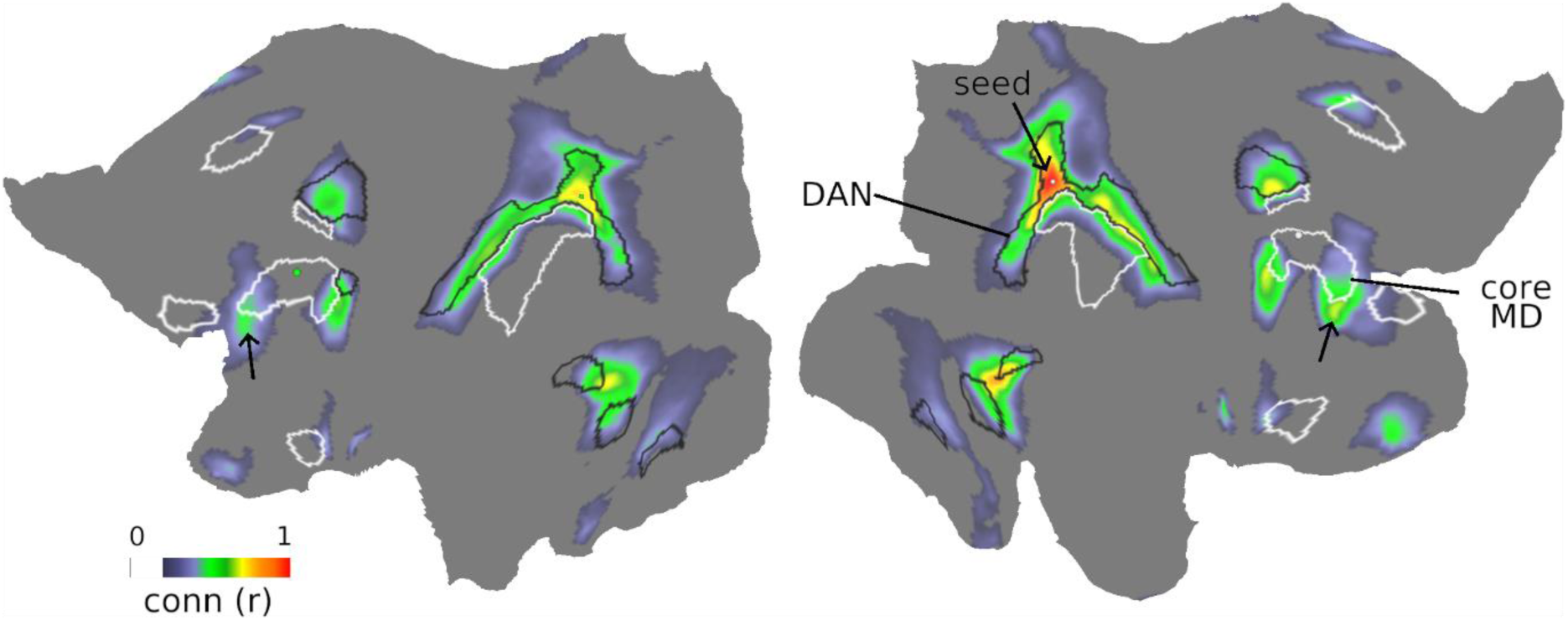
Connectivity (Pearson’s correlation) of a seed in DAN (black borders) shows connectivity to fine-grained regions ventral to the mid-frontal patch of core MD regions (white borders). Correlations are thresholded at 0.2. Correlations are average of the 210 validation HCP subjects (Glasser, Coalson, et al. 2016). Data available at: http://balsa.wustl.edu/qxPk9

**Supplementary Figure 5.**
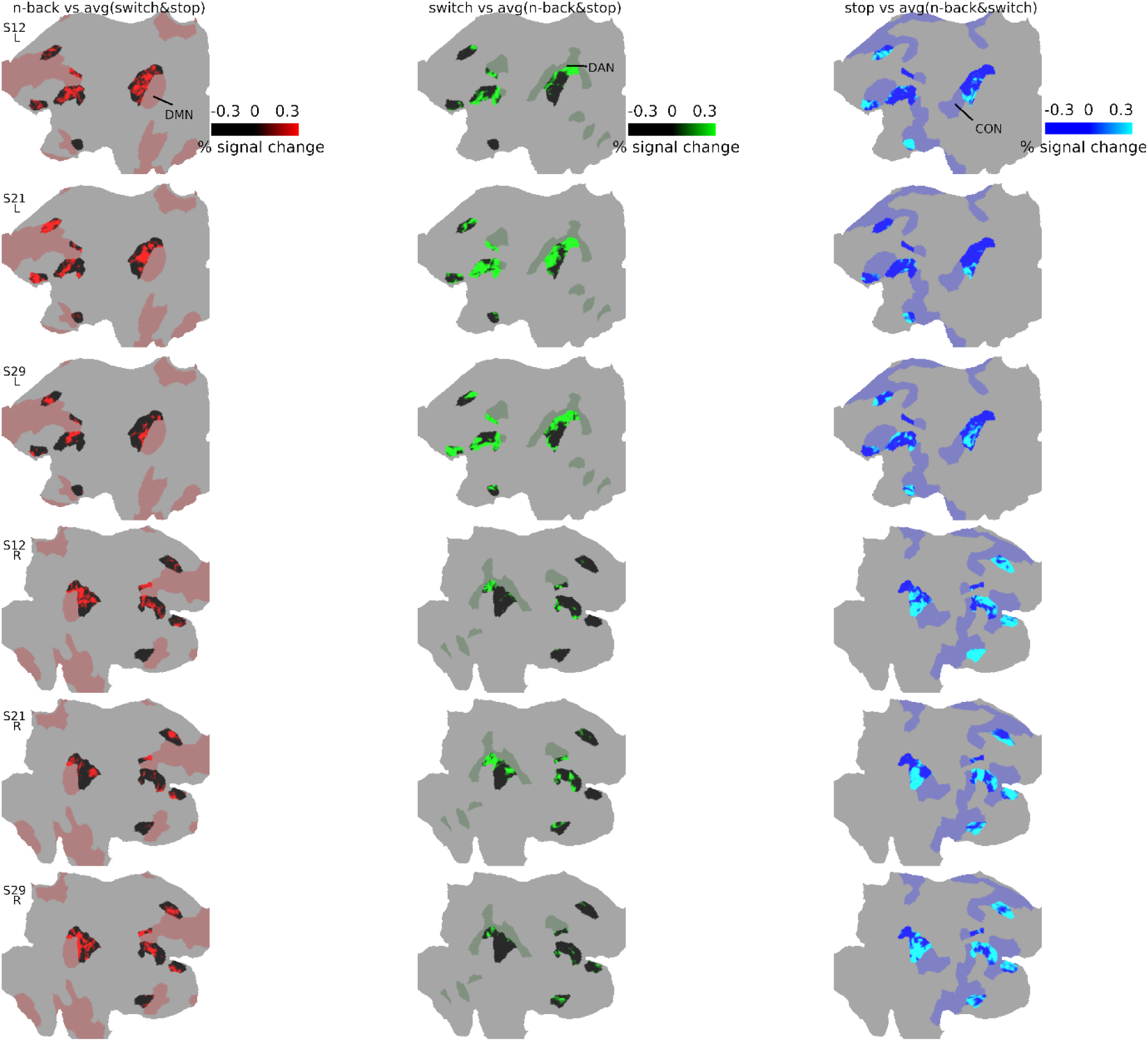
Three example subject activations of each executive task minus the average of the two other tasks. First column shows n-back activations in red surrounded by DMN (faded red). Second column shows switch activations in bright green surrounded by DAN (faded green). Third column shows stop activations in cyan surrounded by CON (faded blue). Data available at: http://balsa.wustl.edu/L7qML

**Supplementary Figure 6.**
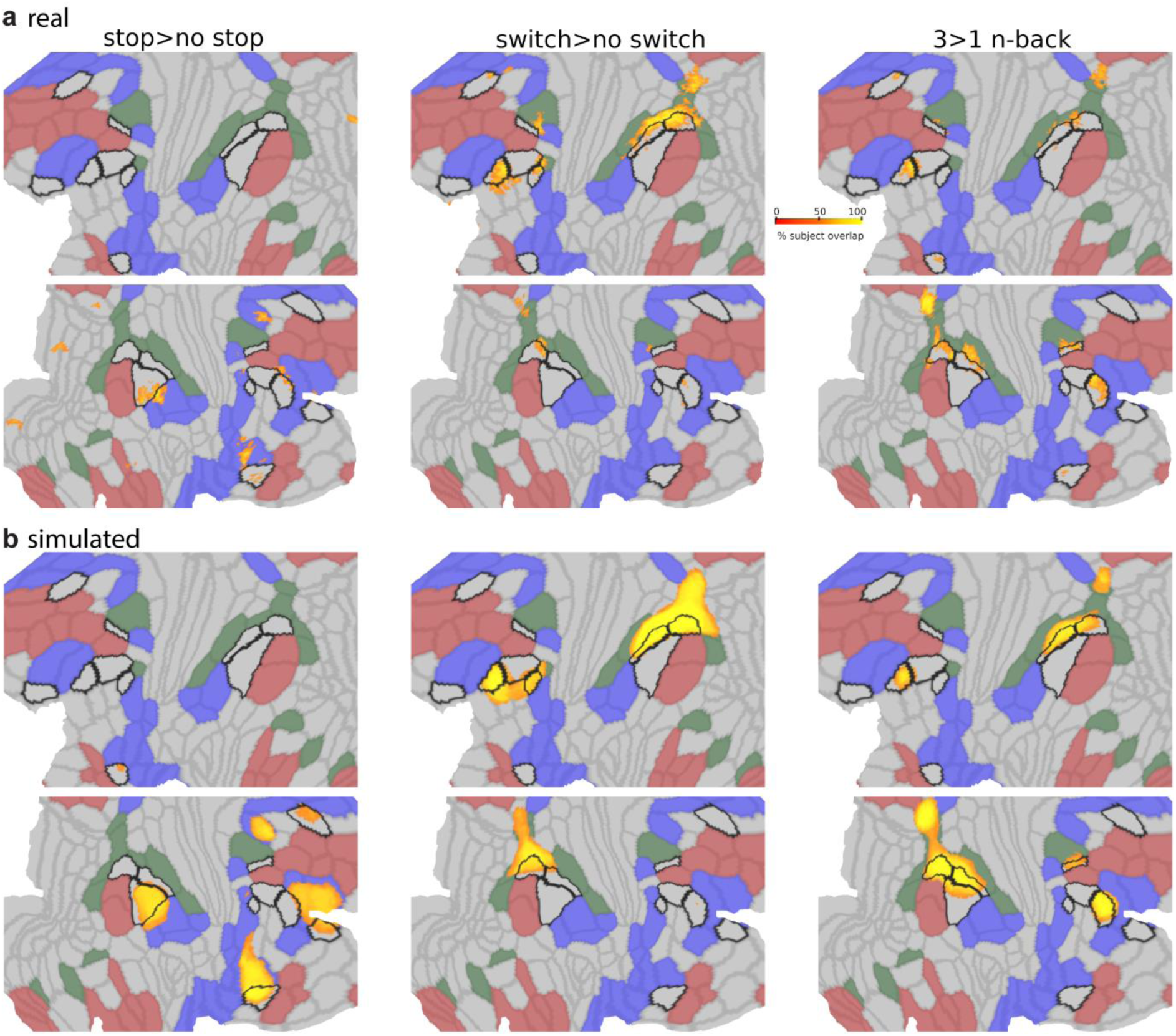
Subject overlap maps for the top 5% vertices in real (top row) and 12 mm smoothed simulated data (bottom row). Data available at: http://balsa.wustl.edu/jNpZl

**Supplementary Figure 7.**
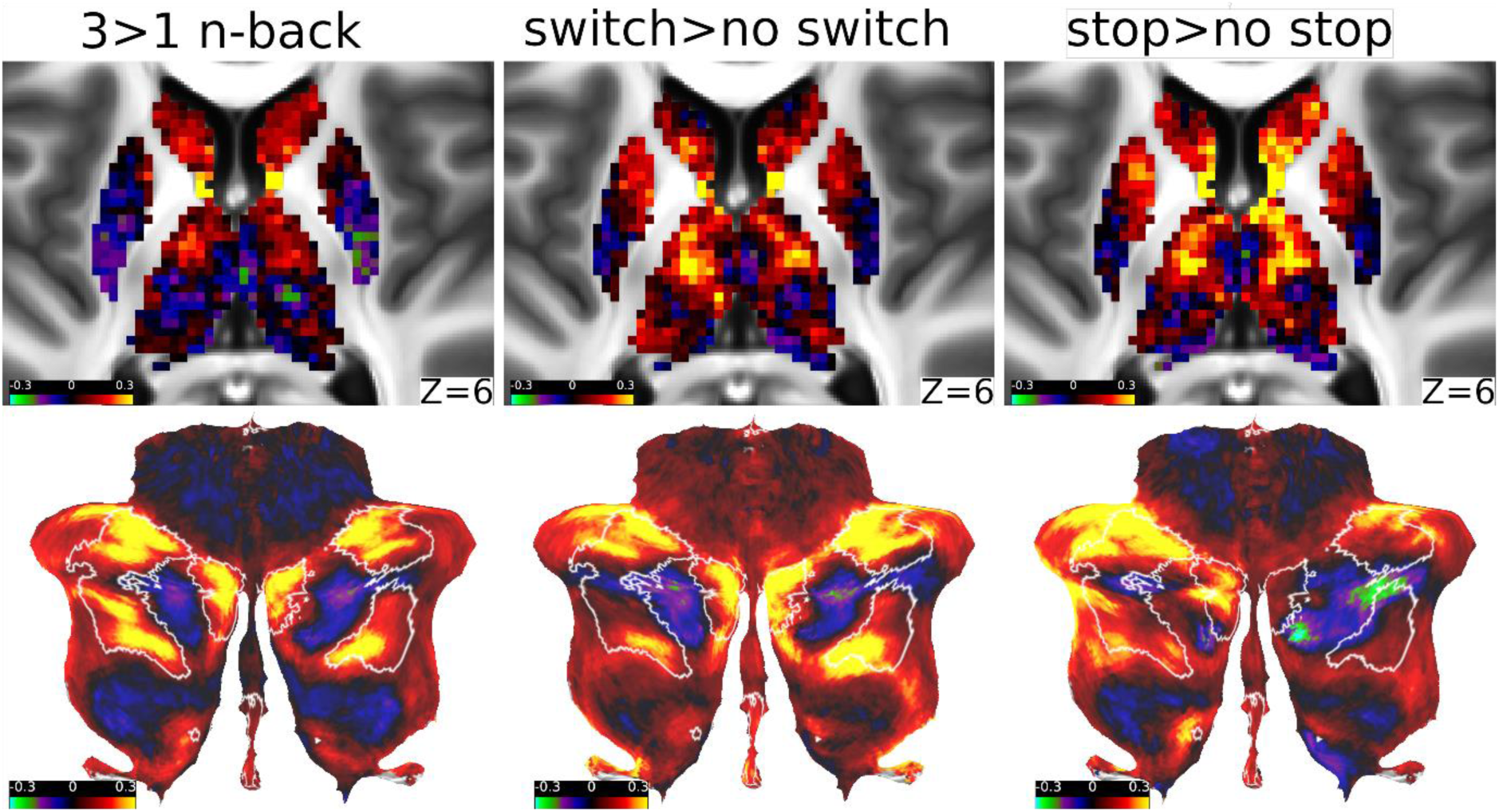
Activations (percent signal change) for each executive contrast in an axial slice of the subcortex (top row) and a flat map of the cerebellum (bottom row). MD areas as defined in (Assem et al. 2020) are surrounded by white borders on the cerebellar surface. Data available at: http://balsa.wustl.edu/w8g71

## References

1. Andrews-Hanna JR, Reidler JS, Sepulcre J, Poulin R, Buckner RL. 2010. Functional-Anatomic Fractionation of the Brain’s Default Network. Neuron. 65:550–562.

2. Apšvalka D, Ferreira CS, Schmitz TW, Rowe JB, Anderson MC. 2022. Dynamic targeting enables domain-general inhibitory control over action and thought by the prefrontal cortex. Nat Commun. 13.

3. Aron AR, Robbins TW, Poldrack RA. 2014. Inhibition and the right inferior frontal cortex: One decade on. Trends Cogn Sci.

4. Arvin O, Keller R, Mefford I, Adams RN. 1978. Lateralization of norepinephrine in human thalamus. Science (80-). 200:1411–1413.

5. Assem M, Glasser MF, Van Essen DC, Duncan J. 2020. A Domain-General Cognitive Core Defined in Multimodally Parcellated Human Cortex. Cereb Cortex. 30:4361– 4380.

6. Assem M, Shashidhara S, Glasser MF, Duncan J. 2022. Precise Topology of Adjacent Domain-General and Sensory-Biased Regions in the Human Brain. Cereb Cortex. 32:2521–2537.

7. Baddeley AD, Hitch G. 1974. Working memory. Psychol Learn Motiv - Adv Res Theory. 8:47–89.

8. Barch DM, Burgess GC, Harms MP, Petersen SE, Schlaggar BL, Corbetta M, Glasser MF, Curtiss S, Dixit S, Feldt C, Nolan D, Bryant E, Hartley T, Footer O, Bjork JM, Poldrack R, Smith S, Johansen-Berg H, Snyder AZ, Van Essen DC. 2013. Function in the human connectome: Task-fMRI and individual differences in behavior. Neuroimage. 80:169–189.

9. Blank IA, Kanwisher N, Fedorenko E. 2014. A functional dissociation between language and multiple-demand systems revealed in patterns of BOLD signal fluctuations. J Neurophysiol. 1105–1118.

10. Braga RM, Van Dijk KRA, Polimeni JR, Eldaief MC, Buckner RL. 2019. Parallel distributed networks resolved at high resolution reveal close juxtaposition of distinct regions. J Neurophysiol. 121:1513–1534.

11. Braver TS. 2012. The variable nature of cognitive control: a dual mechanisms framework. Trends Cogn Sci. 16:106–113.

12. Braver TS, Kizhner A, Tang R, Freund MC, Etzel JA. 2021. The Dual Mechanisms of Cognitive Control Project. J Cogn Neurosci. 1–26.

13. Braver TS, Reynolds JR, Donaldson DI. 2003. Neural Mechanisms of Transient and Sustained Cognitive Control during Task Switching. Neuron. 39:713–726.

14. Coalson TS, Essen DC Van, Glasser MF. 2018. The impact of traditional neuroimaging methods on the spatial localization of cortical areas. Proc Natl Acad Sci. 115:E6356–E6365.

15. Cocuzza C V., Ito T, Schultz D, Bassett DS, Cole MW. 2020. Flexible Coordinator and Switcher Hubs for Adaptive Task Control. J Neurosci. 40:6949–6968.

16. Collette F, Hogge M, Salmon E, Van der Linden M. 2006. Exploration of the neural substrates of executive functioning by functional neuroimaging. Neuroscience. 139:209–221.

17. Collette F, Van der Linden M, Laureys S, Delfiore G, Degueldre C, Luxen A, Salmon E. 2005. Exploring the unity and diversity of the neural substrates of executive functioning. Hum Brain Mapp. 25:409–423.

18. Corbetta M, Shulman GL. 2002. Control of goal-directed and stimulus-driven attention in the brain. Nat Rev Neurosci. 3:201–215.

19. Crone EA, Wendelken C, Donohue SE, Bunge SA. 2006. Neural Evidence for Dissociable Components of Task-switching. Cereb Cortex. 16:475–486.

20. Diamond A. 2013. Executive Functions. Annu Rev Psychol. 64:135–168.

21. Diedrichsen J, Zotow E. 2015. Surface-based display of volume-averaged cerebellar imaging data. PLoS One. 10:1–18.

22. Dodds CM, Morein-Zamir S, Robbins TW. 2011. Dissociating Inhibition, Attention, and Response Control in the Frontoparietal Network Using Functional Magnetic Resonance Imaging. Cereb Cortex. 21:1155–1165.

23. Dosenbach NUF, Visscher KM, Palmer ED, Miezin FM, Wenger KK, Kang HC, Burgund ED, Grimes AL, Schlaggar BL, Petersen SE. 2006. A Core System for the Implementation of Task Sets. Neuron. 50:799–812.

24. Duncan J. 2010. The multiple-demand (MD) system of the primate brain: mental programs for intelligent behaviour. Trends Cogn Sci. 14:172–179.

25. Duncan J, Assem M, Shashidhara S. 2020. Integrated Intelligence from Distributed Brain Activity. Trends Cogn Sci. 24:838–852.

26. Duncan J, Emslie H, Williams P, Johnson R, Freer C. 1996. Intelligence and the Frontal Lobe: The Organization of Goal-Directed Behavior. Cogn Psychol. 30:257–303.

27. Eisenreich BR, Akaishi R, Hayden BY. 2017. Control without Controllers: Toward a Distributed Neuroscience of Executive Control. J Cogn Neurosci. 29:1684–1698.

28. Fedorenko E, Duncan J, Kanwisher N. 2012. Language-selective and domain-general regions lie side by side within Broca’s area. Curr Biol. 22:2059–2062.

29. Fedorenko E, Duncan J, Kanwisher N. 2013. Broad domain generality in focal regions of frontal and parietal cortex. Proc Natl Acad Sci. 110:16616–16621.

30. Friedman NP, Miyake A. 2017. Unity and diversity of executive functions: Individual differences as a window on cognitive structure. Cortex.

31. Friedman NP, Robbins TW. 2022. The role of prefrontal cortex in cognitive control and executive function. Neuropsychopharmacology.

32. Glasser MF, Coalson TS, Bijsterbosch JD, Harrison SJ, Harms MP, Anticevic A, Van Essen DC, Smith SM. 2018. Using temporal ICA to selectively remove global noise while preserving global signal in functional MRI data. Neuroimage.

33. Glasser MF, Coalson TS, Bijsterbosch JD, Harrison SJ, Harms MP, Anticevic A, Van Essen DC, Smith SM. 2019. Classification of temporal ICA components for separating global noise from fMRI data: Reply to Power. Neuroimage. 197:435– 438.

34. Glasser MF, Coalson TS, Robinson EC, Hacker CD, Harwell J, Yacoub E, Ugurbil K, Andersson J, Beckmann CF, Jenkinson M, Smith SM, Van Essen DC. 2016. A multi-modal parcellation of human cerebral cortex. Nature. 536:171–178.

35. Glasser MF, Smith SM, Marcus DS, Andersson JLR, Auerbach EJ, Behrens TEJ, Coalson TS, Harms MP, Jenkinson M, Moeller S, Robinson EC, Sotiropoulos SN, Xu J, Yacoub E, Ugurbil K, Van Essen DC. 2016. The Human Connectome Project’s neuroimaging approach. Nat Neurosci. 19:1175–1187.

36. Glasser MF, Sotiropoulos SN, Wilson JA, Coalson TS, Fischl B, Andersson JL, Xu J, Jbabdi S, Webster M, Polimeni JR, Van Essen DC, Jenkinson M. 2013. The minimal preprocessing pipelines for the Human Connectome Project. Neuroimage. 80:105–124.

37. Hampshire A, Highfield RR, Parkin BL, Owen AM. 2012. Fractionating Human Intelligence. Neuron. 76:1225–1237.

38. He L, Zhuang K, Chen Q, Wei D, Chen X, Fan J, Qiu J. 2021. Unity and diversity of neural representation in executive functions. J Exp Psychol Gen. 150:2193–2207.

39. Isherwood SJS, Keuken MC, Bazin PL, Forstmann BU. 2021. Cortical and subcortical contributions to interference resolution and inhibition – An fMRI ALE meta-analysis. Neurosci Biobehav Rev.

40. Ji JL, Spronk M, Kulkarni K, Repovš G, Anticevic A, Cole MW. 2019. Mapping the human brain’s cortical-subcortical functional network organization. Neuroimage. 185:35–57.

41. Karr JE, Areshenkoff CN, Rast P, Hofer SM, Iverson GL, Garcia-Barrera MA. 2018. The unity and diversity of executive functions: A systematic review and re-analysis of latent variable studies. Psychol Bull. 144:1147–1185.

42. Lemire-Rodger S, Lam J, Viviano JD, Stevens WD, Spreng RN, Turner GR. 2019. Inhibit, switch, and update: A within-subject fMRI investigation of executive control. Neuropsychologia. 132:107134.

43. Meuwissen AS, Anderson JE, Zelazo PD. 2017. The creation and validation of the Developmental Emotional Faces Stimulus Set. Behav Res Methods. 49:960–966.

44. Michalka SW, Kong L, Rosen ML, Shinn-Cunningham BG, Somers DC. 2015. Short-Term Memory for Space and Time Flexibly Recruit Complementary Sensory-Biased Frontal Lobe Attention Networks. Neuron. 87:882–892.

45. Miyake A, Friedman NP, Emerson MJ, Witzki AH, Howerter A, Wager TD. 2000. The Unity and Diversity of Executive Functions and Their Contributions to Complex “Frontal Lobe” Tasks: A Latent Variable Analysis. Cogn Psychol. 41:49–100.

46. Murphy C, Jefferies E, Rueschemeyer S-A, Sormaz M, Wang H, Margulies DS, Smallwood J. 2018. Distant from input: Evidence of regions within the default mode network supporting perceptually-decoupled and conceptually-guided cognition. Neuroimage. 171:393–401.

47. Nee DE. 2021. Integrative frontal-parietal dynamics supporting cognitive control. Elife. 10.

48. Nee DE, Brown JW, Askren MK, Berman MG, Demiralp E, Krawitz A, Jonides J. 2013. A Meta-analysis of Executive Components of Working Memory. Cereb Cortex. 23:264–282.

49. Niendam TA, Laird AR, Ray KL, Dean YM, Glahn DC, Carter CS. 2012. Meta-analytic evidence for a superordinate cognitive control network subserving diverse executive functions. Cogn Affect Behav Neurosci. 12:241–268.

50. Panichello MF, Buschman TJ. 2021. Shared mechanisms underlie the control of working memory and attention. Nature. 592:601–605.

51. Power JD, Cohen AL, Nelson SM, Wig GS, Barnes KA, Church JA, Vogel AC, Laumann TO, Miezin FM, Schlaggar BL, Petersen SE. 2011. Functional Network Organization of the Human Brain. Neuron. 72:665–678.

52. Power JD, Schlaggar BL, Lessov-Schlaggar CN, Petersen SE. 2013. Evidence for hubs in human functional brain networks. Neuron. 79:798–813.

53. Reineberg AE, Banich MT, Wager TD, Friedman NP. 2022. Context-specific activations are a hallmark of the neural basis of individual differences in general executive function. Neuroimage. 249:118845.

54. Robinson EC, Garcia K, Glasser MF, Chen Z, Coalson TS, Makropoulos A, Bozek J, Wright R, Schuh A, Webster M, Hutter J, Price A, Cordero Grande L, Hughes E, Tusor N, Bayly P V., Van Essen DC, Smith SM, Edwards AD, Hajnal J, Jenkinson M, Glocker B, Rueckert D. 2018. Multimodal surface matching with higher-order smoothness constraints. Neuroimage. 167:453–465.

55. Robinson EC, Jbabdi S, Glasser MF, Andersson J, Burgess GC, Harms MP, Smith SM, Van Essen DC, Jenkinson M. 2014. MSM: A new flexible framework for multimodal surface matching. Neuroimage. 100:414–426.

56. Roca M, Parr A, Thompson R, Woolgar A, Torralva T, Antoun N, Manes F, Duncan J. 2010. Executive function and fluid intelligence after frontal lobe lesions. Brain. 133:234–247.

57. Rugg MD, Vilberg KL. 2013. Brain networks underlying episodic memory retrieval. Curr Opin Neurobiol.

58. Salimi-Khorshidi G, Douaud G, Beckmann CF, Glasser MF, Griffanti L, Smith SM. 2014. Automatic denoising of functional MRI data: Combining independent component analysis and hierarchical fusion of classifiers. Neuroimage. 90:449– 468.

59. Saylik R, Williams AL, Murphy RA, Szameitat AJ. 2022. Characterising the unity and diversity of executive functions in a within-subject fMRI study. Sci Rep. 12:8182.

60. Sebastian A, Konken AM, Schaum M, Lieb K, Tüscher O, Jung P. 2021. Surprise: Unexpected action execution and unexpected inhibition recruit the same fronto-basal-ganglia network. J Neurosci. 41:2447–2456.

61. Seghezzi S, Haggard P. 2022. Volition and “free will.” psyarxiv.

62. Shao X, Krieger-Redwood K, Zhang M, Hoffman P, Lanzoni L, Leech R, Smallwood J, Jefferies E. 2023. Distinctive and complementary roles of default mode network subsystems in semantic cognition. bioRxiv. 2023.10.03.560166.

63. Shashidhara S, Spronkers FS, Erez Y. 2020. Individual-subject Functional Localization Increases Univariate Activation but Not Multivariate Pattern Discriminability in the “Multiple-demand” Frontoparietal Network. J Cogn Neurosci. 32:1348–1368.

64. Si R, Rowe JB, Zhang J. 2021. Functional localization and categorization of intentional decisions in humans: A meta-analysis of brain imaging studies. Neuroimage. 242. Spearman C. 1904. “General Intelligence,” Objectively Determined and Measured. Am J Psychol. 15:201.

65. Stuss DT. 2011. Functions of the Frontal Lobes: Relation to Executive Functions. J Int Neuropsychol Soc. 17:759–765.

66. Stuss DT, Alexander MP. 2000. Executive functions and the frontal lobes: a conceptual view. Psychol Res. 63:289–298.

67. Suda A, Osada T, Ogawa A, Tanaka M, Kamagata K, Aoki S, Hattori N, Konishi S. 2020. Functional organization for response inhibition in the right inferior frontal cortex of individual human brains. Cereb Cortex. 30:6325–6335.

68. Swann NC, Cai W, Conner CR, Pieters TA, Claffey MP, George JS, Aron AR, Tandon N. 2012. Roles for the pre-supplementary motor area and the right inferior frontal gyrus in stopping action: Electrophysiological responses and functional and structural connectivity. Neuroimage. 59:2860–2870.

69. Tavor I, Jones OP, Mars RB, Smith SM, Behrens TE, Jbabdi S. 2016. Task-free MRI predicts individual differences in brain activity during task performance. Science (80-). 352:216–220.

70. Tsumura K, Aoki R, Takeda M, Nakahara K, Jimura K. 2021. Cross-hemispheric complementary prefrontal mechanisms during task switching under perceptual uncertainty. J Neurosci. 41:2197–2213.

71. Wager TD, Sylvester CYC, Lacey SC, Nee DE, Franklin M, Jonides J. 2005. Common and unique components of response inhibition revealed by fMRI. Neuroimage. 27:323–340.

72. Warren DE, Power JD, Bruss J, Denburg NL, Waldron EJ, Sun H, Petersen SE, Tranel D. 2014. Network measures predict neuropsychological outcome after brain injury. Proc Natl Acad Sci. 111:14247–14252.

73. Waskom ML, Wagner AD. 2017. Distributed representation of context by intrinsic subnetworks in prefrontal cortex. Proc Natl Acad Sci U S A. 114:2030–2035.

74. Wen T, Mitchell DJ, Duncan J. 2020. The Functional Convergence and Heterogeneity of Social, Episodic, and Self-Referential Thought in the Default Mode Network. Cereb Cortex. 30:5915–5929.

75. Woolgar A, Duncan J, Manes F, Fedorenko E. 2018. Fluid intelligence is supported by the multiple-demand system not the language system. Nat Hum Behav. 2:200– 204.

76. Woolgar A, Parr A, Cusack R, Thompson R, Nimmo-Smith I, Torralva T, Roca M, Antoun N, Manes F, Duncan J. 2010. Fluid intelligence loss linked to restricted regions of damage within frontal and parietal cortex. Proc Natl Acad Sci U S A. 107:14899–14902.

77. Woolrich MW, Ripley BD, Brady M, Smith SM. 2001. Temporal Autocorrelation in Univariate Linear Modeling of FMRI Data. Neuroimage. 14:1370–1386.

78. Yeo BTT, Krienen FM, Eickhoff SB, Yaakub SN, Fox PT, Buckner RL, Asplund CL, Chee MWL. 2015. Functional Specialization and Flexibility in Human Association Cortex. Cereb Cortex. 25:3654–3672.

79. Yeo BTT, Krienen FM, Sepulcre J, Sabuncu MR, Lashkari D, Hollinshead M, Roffman JL, Smoller JW, Zöllei L, Polimeni JR, Fischl B, Liu H, Buckner RL. 2011. The organization of the human cerebral cortex estimated by intrinsic functional connectivity. J Neurophysiol. 106:1125–1165.

